# Converting antimicrobial into targeting peptides reveals key features governing protein import into mitochondria and chloroplasts

**DOI:** 10.1101/2021.12.03.471120

**Authors:** Oliver D Caspari, Clotilde Garrido, Chris O Law, Yves Choquet, Francis-André Wollman, Ingrid Lafontaine

## Abstract

We asked what peptide features govern targeting to the mitochondria versus the chloroplast using antimicrobial peptides as a starting point. This approach was inspired by the endosymbiotic hypothesis that organelle-targeting peptides derive from antimicrobial amphipathic peptides delivered by the host cell, to which organelle progenitors became resistant. To explore the molecular changes required to convert antimicrobial into targeting peptides, we expressed a set of 13 antimicrobial peptides in *Chlamydomonas reinhardtii*. Peptides were systematically modified to test distinctive features of mitochondrial and chloroplast targeting peptides, and we assessed their targeting potential by following the intracellular localization and maturation of a Venus fluorescent reporter used as cargo protein. Mitochondrial targeting can be achieved by some unmodified antimicrobial peptide sequences. Targeting to both organelles is improved by replacing Lysines with Arginines. Chloroplast targeting is enabled by the presence of flanking unstructured sequences, additional constraints consistent with chloroplast endosymbiosis having occurred in a cell that already contained mitochondria. If indeed targeting peptides evolved from antimicrobial peptides, required modifications imply a temporal evolutionary scenario with an early exchange of cationic residues, and a late acquisition of chloroplast specific motifs.

## Introduction

Mitochondria and chloroplasts arose through endosymbiosis and retain their own genomes, yet the vast majority of organellar proteins is encoded in the nucleus, translated in the cytoplasm and imported into the organelles (Wiedemann and Pfanner, 2017; Chotewutmontri et al., 2017). N-terminal targeting peptides (TPs) that are cleaved off upon import provide the information on targeting, although their primary structures are very diverse (Bruce, 2001). Yet chloroplast transit peptides (cTP) and mitochondrial presequences (mTP) have very similar physico-chemical properties making reliable differentiation often challenging. A multitude of studies has found sequence elements contributing to specificity determination, and prediction programs have been improving (Tardif et al., 2012; Armenteros et al., 2019), yet a mechanistic understanding of how targeting information is encoded has remained elusive.

Here, we use antimicrobial peptides (AMP) as an original chassis, to test the contribution of key TP features towards targeting efficiency and specificity. Part of the innate immune system, AMPs are produced by virtually all types of cells in a bid to kill or control microbial adversaries (Joo et al., 2016; Lazzaro et al., 2020). AMPs have recently been proposed to be at the evolutionary origin of TPs (Wollman, 2016; Caspari and Lafontaine, 2021). The proposed evolutionary scenario posits that early during endosymbiotic organellogenesis of first the mitochondrion and later the chloroplast, the host cell used AMPs to attack the bacterial proto-endosymbiont. A bacterial resistance mechanism, whereby the AMP is imported into the bacterial cytoplasm, would have generated a pathway for host proteins to reach the bacterial cytosol – a plausible first step in the evolution of a protein import machinery. Cationic, Helical-Amphipathic Ribosomally-produced AMPs (HA-RAMPs) and TPs share key physico-chemical properties and they were shown, in some instances, to retain cross-functionality (Garrido et al., 2020): several TPs showed antimicrobial activity and selected HA-RAMPs, fused to a cleavage-site containing TP element, were shown to promote the import of a Venus fluorescent protein into either the mitochondria or the chloroplast of the model green alga *Chlamydomonas reinhardtii*.

The main part of mTPs and the central element of cTPs most closely resemble HA-RAMPs on account of a shared cationic, amphipathic helical structure that often encompasses the entire length of HA-RAMPs (Caspari and Lafontaine, 2021) and mTPs (von Heijne, 1986; von Heijne et al., 1989). Although plant cTPs have been described as unstructured (von Heijne and Nishikawa, 1991), algal cTPs more closely resemble mTPs in being helical (Franzén et al., 1990). Helices were observed by NMR in membrane-mimetic environments in both algal and plant cTPs (Lancelin et al., 1996; Bruce, 1998; Krimm et al., 1999; Wienk et al., 2000; Bruce, 2001), and signatures of amphipathic helices can be detected in a majority of cTPs (Garrido et al., 2020).

In addition to the helices, both mTPs and cTPs contain recognition sites at the C-terminus where processing peptidases in the mitochondrial matrix (MPP) and the chloroplast stroma (SPP) cleave off the cargo protein (Teixeira and Glaser, 2013). These recognition sites encompass some 10 residues upstream of the cleavage site and are structurally distinct from the rest of the TPs, even showing a weak sequence conservation (von Heijne et al., 1989; Tardif et al., 2012; Köhler et al., 2015). While targeting information is usually contained within mTP sequences upstream of the cleavage site, targeting by cTPs shorter than ca. 60 amino acids often requires downstream unstructured sequence stretches in the N-terminal domain of the mature protein (Bionda et al., 2010; Caspari, 2022). Besides the amphipathic helical module and the C-terminal cleavage module shared between mTPs and cTPs, it has been argued that distinct features at their N-termini conferred organelle specificity to each set of targeting peptides (von Heijne et al., 1989; Ivey and Bruce, 2000; Ivey et al., 2000; Bhushan et al., 2006; Chotewutmontri et al., 2012; Chotewutmontri and Bruce, 2015; Köhler et al., 2015; Chotewutmontri et al., 2017; Lee et al., 2019).

In this study, we systematically introduced modifications into a diversity of HA-RAMPs in a bid to generate targeting to the mitochondria or chloroplast in Chlamydomonas. This dataset provides new insights into how different TP elements contribute to differential targeting and to the efficiency of protein import. Being similar in physico-chemical properties to TPs (Garrido et al., 2020), HA-RAMPs provide a privileged vantage point from which to study sequence elements that govern targeting. In our choice of HA-RAMPs, we aimed to reflect the diversity of available sequences by choosing representatives of different HA-RAMP families based on similarity with TPs. We show that some of our 13 HA-RAMPs natively contain TP-like sequence elements, with some HA-RAMPs being prone to chloroplast targeting while others show a preference for the mitochondria. Furthermore, we provide evidence for a critical functional difference in cationic residues with Lysine (K) being used in HA-RAMPs and Arginine (R) in TPs.

## Results

### HA-RAMPs display varying degrees of similarity to TPs

Figure 1 shows the major sequence features of the 13 HA-RAMPs used in the present study (Fig. 1a) together with those of a typical cTP and a typical mTP (Fig. 1b). On the right-hand side, these peptides are represented according to their proportion of a-helical amphipathic structure. The HA-RAMPs Brevinin-2ISb (B2I), Magainin 2 (MII), Ranatuerin-2G (R2G), Dermaseptin S4 (DS4), Dermadistinctin-M (DDM), Brevinin-1E (B1E), Cecropin-P3 (CP3), Sarcotoxin-1D (S1D), Esculentin-1SEA (E1S), Leucocin A (LCA), SI Moricin (SIM), Bacillocin 1580 (B15), and Enterocin HF (EHF) were chosen such that different AMP families would be represented with varying proximity to TPs (Table S1) and thus a range of physico-chemical properties be explored (Fig. S1). As negative controls, two peptides that lack predicted amphipathic helices were chosen from among randomly generated sequences (Fig. 1c).

**Fig. 1.**
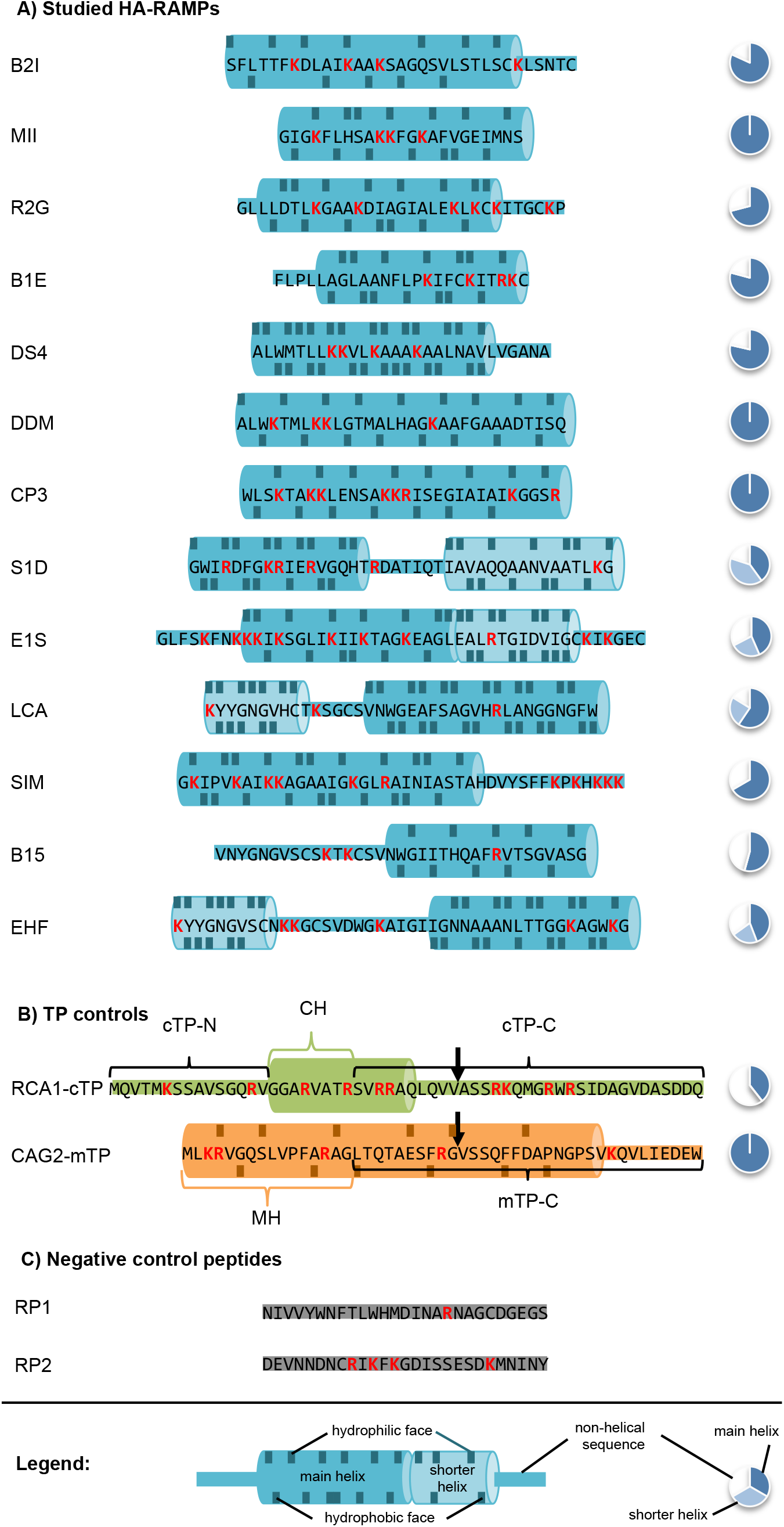
Peptide sequences under study. Amino acid sequences are shown using the one-letter code. Positively charged residues are highlighted in red. The fraction of the sequence predicted to fold into amphipathic helices (see Methods) is provided by a pie-chart to the right of the sequence; for TPs in (**B**) this was calculated up to the cleavage site indicated by a downward arrow. Predicted amphipathic helices are highlighted using a cylinder cartoon, with residues contributing to the hydrophilic/hydrophobic face indicated on the top/bottom. No helix could be predicted within RCA1-cTP, and thus the indicated helix is taken from a published NMR-structure obtained in membrane-mimetic conditions (Krimm et al., 1999). Note that the two helices of E1S are at an angle to each other and therefore cannot form a single continuous amphipathic helix. Abbreviations: (**A**) B2I: Brevinin-2ISb, R2G: Ranatuerin-2G, MII: Magainin 2, B1E: Brevinin-1E, DS4: Dermaseptin S4, DDM: Dermadistinctin-M, CP3: Cecropin-P3, S1D: Sarcotoxin-1D, E1S: Esculentin-1SEA, LCA: Leucocin-A, SIM: SI Moricin, B15: Bacillocin 1580, EHF: Enterocin HF; (**B**) TP – targeting peptide, cTP: chloroplast TP, mTP: mitochondrial TP, RCA1: Rubisco activase, CAG2: γ-carbonic anhydrase, cTP-N: cTP N-terminal element, CH: cTP helix, MH: mTP helix, cTP-C: cTP C-terminal element, mTP-C: mTP C–terminal element; (**C**) RP: random peptide.

To explore targeting, the 13 HA-RAMPs were systematically modified (Fig. 2), notably by adding TP elements from the N- and/or C-terminal domains of Chlamydomonas Rubisco activase (RCA1) cTP, or a C-terminal domain of similar length from mitochondrial γ-carbonic anhydrase 2 (CAG2) mTP (Fig. 1b, Fig. 2). In a bid to keep peptidase recognition sites intact, C-terminal elements were designed to start 10 residues upstream of the cleavage site, even though this slightly truncates TP helices. Peptides carrying the 15 amino-acid long cTP N-terminus (cTP-N) will be denoted ^c^P. Similarly, peptides with C-terminal elements from RCA1-cTP (cTP-C) or CAG2-mTP (mTP-C) will be denoted P^c^ or P^m^ respectively. Peptide variants are used to drive the subcellular localization of a Venus fluorescent reporter protein, which was assessed using fluorescence microscopy; in Fig. 2 an executive overview of observed Venus localization is presented for all constructs. We previously validated fluorescence localization biochemically in a small number of strains by showing that the Venus-FLAG reporter is retained within isolated mitochondria or chloroplasts (Garrido et al., 2020). Here, we use automated image segmentation to validate our subcellular localization assessment (Fig. S2, Supplementary text). Localization was obtained using stable Chlamydomonas expression lines, generated by introducing DNA sequences encoding peptides upstream of the Venus coding sequence in a bicistronic expression vector (Caspari, 2020) using Gibson assembly, with transformation cassettes integrated into the Chlamydomonas nuclear genome at random sites via electroporation. Micrographs for three biological replicates (i.e., independent insertion lines) per construct are displayed in Fig. S4-S21. The reader will be asked to return to Fig. 2 throughout, with Fig. 3 and Fig. 4 providing selected micrographs as examples highlighting particular points of interest.

**Fig. 2.**
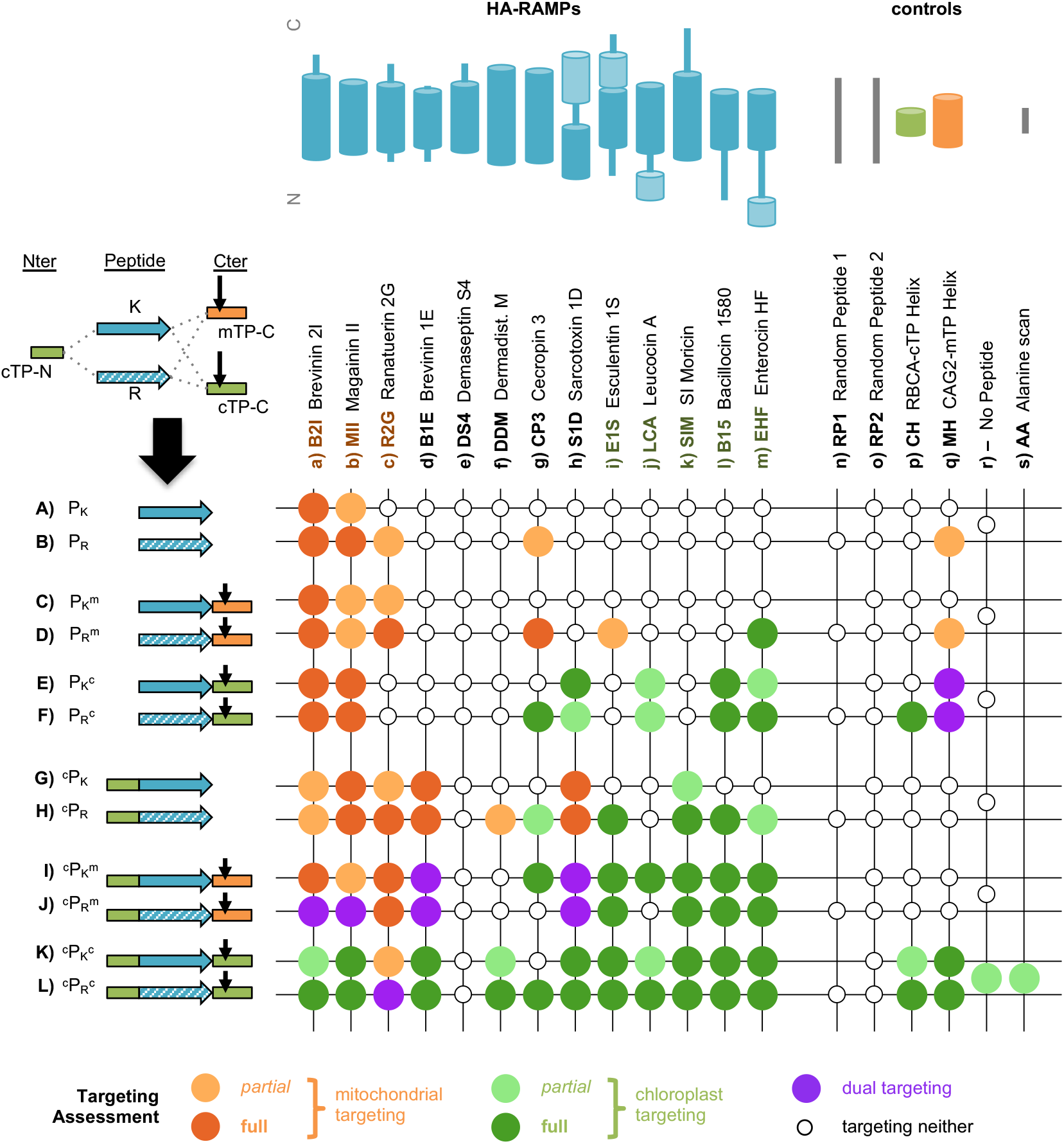
TP modifications enable HA-RAMP targeting. Chimeric constructs were generated by combining TP elements with HA-RAMPs (see Fig. 1 for sequences; in the ^c^AA^c^ construct in column s, all residues of the helical element “CH” within RCA1-cTP are replaced by Alanines). The overview graph shows in each column (**a-s**) one of the peptides with a cartoon indicating the position of the predicted amphipathic helices within the sequence; and in each row (**A-L**) a combination of peptide, K/R modification, addition of cTP-N (the 15 N-terminal residues upstream of the helix in RCA1-cTP) and/or a C-terminal TP element (cTP-C or mTP-C, which include −10 residues upstream and +23 residues downstream of the cleavage site for RCA1-cTP and CAG2-mTP respectively) indicated by a cartoon and the following shorthand: P = peptide, K = contains mostly Lysines, R = contains mostly Arginines, ^m^ = mTP element, ^c^ = cTP element (^c^P = cTP-N, P^c^ = cTP-C). In each case, an overview of observed targeting is provided by a colour code (see figure legend). Images for all constructs are shown in Fig. S3-S20. Note that the present results for B2I_K_^c^, MII_K_^c^, S1D_K_^c^, B15_K_^c^ and EHF_K_^c^ (row E, columns a,b,h,l,m) confirmed our previous report on these strains (Garrido et al., 2020).

**Fig. 3.**
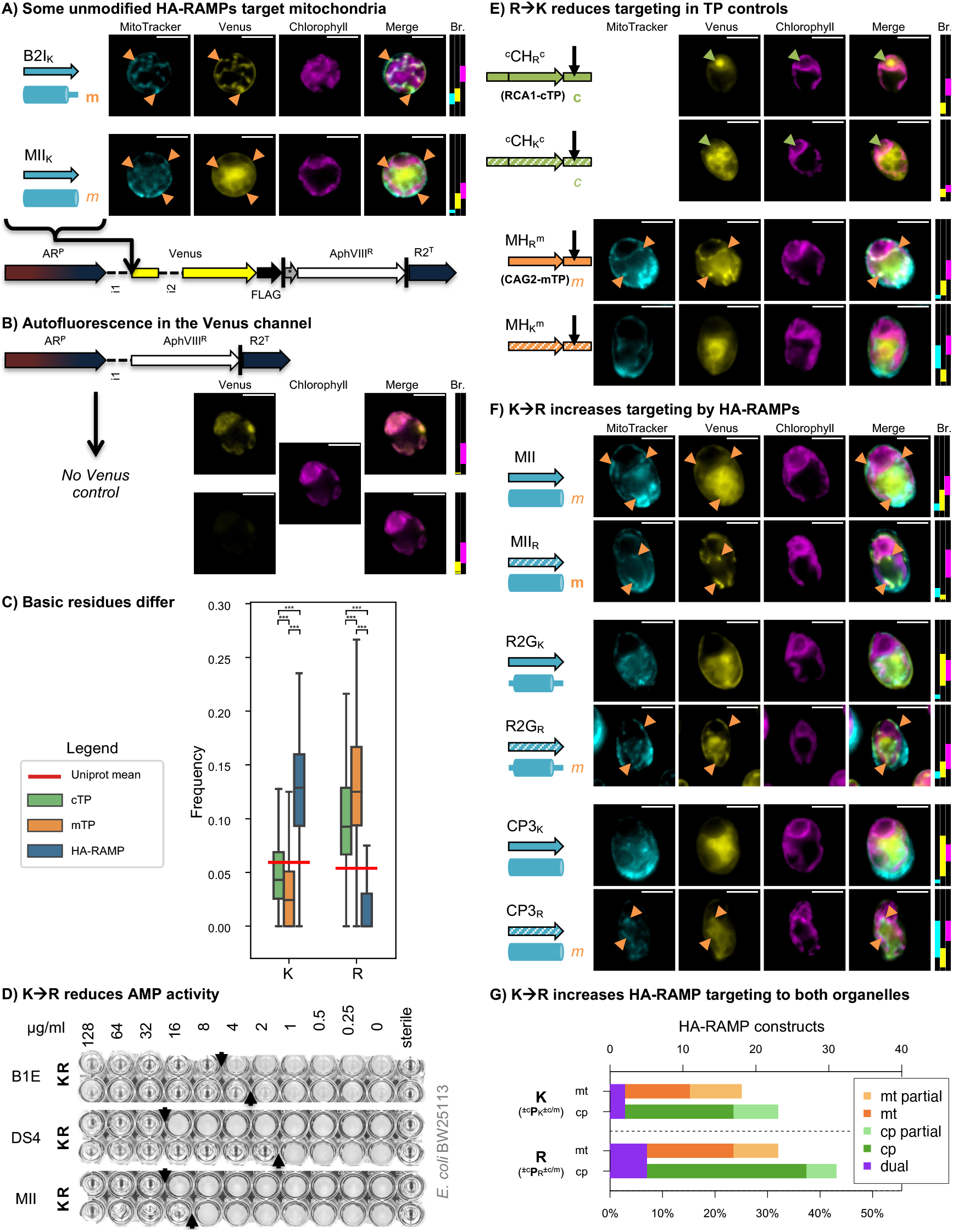
K is for killing, R is for targeting. (**A**) Indicated peptides were inserted upstream of a Venus fluorescent protein reporter in a bicistronic expression system for Chlamydomonas (Caspari, 2020). AR^P^: hybrid *HSP70a-RBCS2* promoter, i1: *RBCS2* intron 1, i2: *RBCS2* intron 2, FLAG: FLAG-tag, |: stop codon, *: bicistronic bridge sequence tagcat, AphVIII^R^: Paromomycin resistance gene, R2^T^: *RBCS2* terminator. Epifluorescence microscopy images of selected examples are shown. False-coloured yellow fluorescence from the Venus channel reports on the subcellular localization of the fluorescent reporter. MitoTracker fluorescence, false-coloured in cyan, indicates the position of mitochondria (although parts of the cell exterior are sometimes also stained), with salient features highlighted with orange arrows to indicate co-localization with the Venus channel. Chlorophyll autofluorescence, shown in magenta, indicates the location of the chloroplast. Scale bars are 5μm. Refer to Fig. 1 for sequences and Fig. S3 and S4 for biological replicates. Where a construct was interpreted as generating reporter localization in mitochondria or chloroplast, this is indicated by an orange ‘m’ or a green ‘c’ respectively, in bold for full targeting or in italics for partial targeting. Brightness (Br.) was adjusted for clarity: fluorescence intensity values were restricted to the range shown for each channel by matching coloured rectangles. Intensity scales to 0 at the bottom of the panel, and to 65535 at the top. (**B**) A ‘no Venus’ control strain, expressing an empty vector, is shown with two different Venus channel brightness settings to visualize chloroplast autofluorescence in the Venus channel. Autofluorescence intensity is typically below 2000, indicated by a black dotted line in Brightness rectangles. Therefore, if Venus channel brightness is adjusted below 2000, autofluorescence originating from the chloroplast may be misinterpreted as Venus located in the chloroplast. **(C)** Lysine (K) and Arginine (R) frequencies for Chlamydomonas TPs and HA-RAMPs are shown as boxplots (centre line: median; box limits: upper and lower quartiles; whiskers: min/max values within 1.5x interquartile range). To give a baseline for comparison, their average across the UNIPROT database is given as red horizontal line. Statistically significant differences are indicated with stars: (Multiple Kruskal-Wallis tests followed by Dunn post-hoc tests, ***p<0.0001). See Fig. S21 for all amino acids. (**D**) Grayscale photos of antimicrobial activity growth assays in the presence of dilutions of three selected antimicrobial peptides against *E. coli* strain BW25113 show reduced activity when natively K-rich sequences (K, left hand columns) are altered by replacing all Ks by Rs (R, right hand columns). (**E,F**) Epifluorescence images of selected chimeric (**E**) TP controls (MH: mTP helical fragment, CH: cTP helical fragment) or (**F**) HA-RAMP constructs are shown as in (A). Note that images shown for MII_K_ in (A) and (F) come from independent insertion lines of the same construct. See Fig. S4, S5, S9, S18, S19 for biological replicates. (**G**) Data for targeting across all HA-RAMP constructs (i.e. including additional modifications) is shown, comparing K and R–bearing peptides (cf. Fig. 2 columns a-m).

**Fig. 4.**
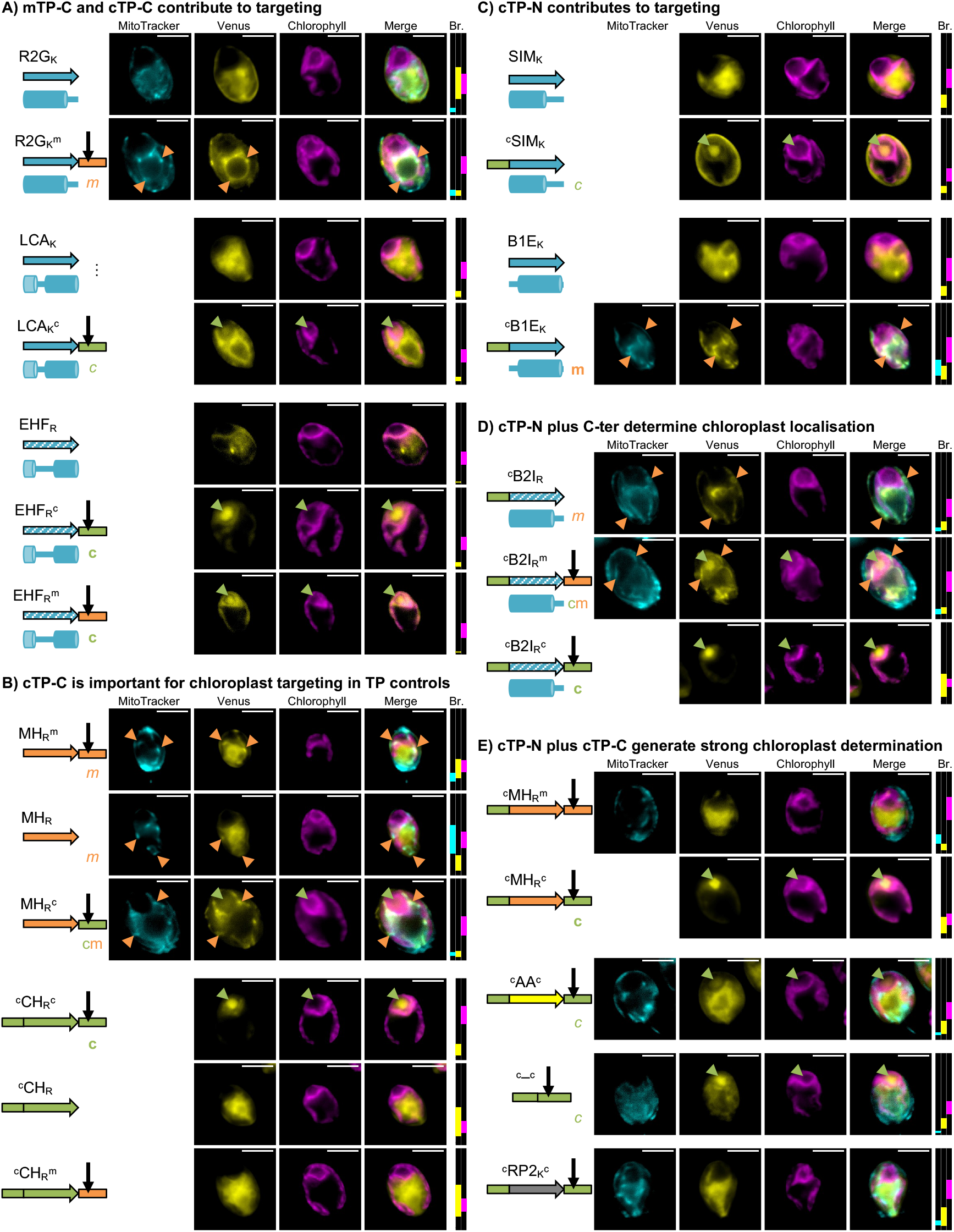
TP N- and C-termini enable chloroplast targeting by HA-RAMPs. Exemplary epifluorescence images are shown as in Fig. 3; same convention as in Fig. 3 for Venus localization. Note that images shown for MH_R_^m^ and ^c^CH_R_^c^ in (B) come from different independent insertion lines of the same constructs as in Fig. 3E. See Fig. S3-S20 for biological replicates and further examples of the same trends.

### Some unmodified HA-RAMPs generate mitochondrial targeting

In the absence of any modifications, B2I and MII were capable of organelle-targeting visible in fluorescence microscopy (Fig. 2 row A, Fig. S3-S15). When equipped with B2I, the fluorescent reporter Venus shows mitochondrial localization (Fig. 3A, Fig. S3): the Venus signal is observed as a characteristic network pattern that co-localizes with MitoTracker fluorescence (cf. Fig. S2A, S4). By contrast, in the case of MII only a fraction of the fluorescence signal in the Venus channel shows co-localization with the MitoTracker dye, signifying that targeting is only partial (Fig. 3A, Fig. S5). Note that in these epifluorescence images, some autofluorescence emanating from the chloroplast is present in the Venus channel (Fig. 3B). In all images, display brightness was adjusted such that Venus localization would be clearly visible, and where possible this included removing autofluorescence. Brightness settings were chosen as indicated next to each image for full transparency. In cases where low Venus accumulation necessitated brightness values low enough for autofluorescence to be visible in the Venus channel, a black dotted line is visible in the brightness display, indicating that some autofluorescence signal co-localizing with chlorophyll should be expected in the Venus channel independent of the genuine Venus localization.

### K/R content contributes to functional divergence between HA-RAMP and TP

Extant HA-RAMPs and organellar TPs display very few differences in their amino-acid content (Supplementary text, Fig. S20). As expected, both are poor in acidic residues (D, E) but enriched in basic residues (K or R). However, their complement in basic residues is markedly different (Fig. 3C): HA-RAMPs are rich in Lysines (K) whereas TPs are rich in Arginines (R).

In order to see whether these contrasting differences in K/R ratio contributed to the functional divergence between HA-RAMPs and TPs, we substituted all instances of K by R in HA-RAMPs and of R by K in TPs. In the rest of the text, the basic amino acid mostly present in a given peptide P are indicated by a subscript as P_R_ or P_K_. HA-RAMPs with K→R transition showed reduced antimicrobial activity, as illustrated on Fig. 3D by the increased minimum inhibitory concentrations for MII, DS4 or B1E.

We used Chlamydomonas Rubisco activase (RCA1) cTP (^C^CH_R_^c^) and mitochondrial γ-carbonic anhydrase 2 (CAG2) mTP (MH_R_^m^) as positive controls for chloroplast and mitochondrial targeting respectively (Fig. 3E, Fig. S18-S19). Note that TP helical fragments, stripped of their N- and C-terminal domains, are denoted as MH for mTP and CH for cTP (as detailed in Fig. 1B). When equipped with RCA1-cTP (^C^CH_R_^c^), Venus shows two chloroplast localization features: a diffuse signal that colocalizes with chlorophyll fluorescence, and a bright spot where there is a drop in chlorophyll fluorescence. Both of these features are genuine markers of chloroplast localization: in Chlamydomonas, the single cup-shaped chloroplast has a reliable morphology, and the dip in chlorophyll fluorescence at the base of the chloroplast is a well-established marker of the pyrenoid, a proteinaceous chloroplast sub-compartment that contains a large majority of Rubisco (Fig. S2B) (Mackinder et al., 2016; Caspari et al., 2017; Mackinder et al., 2017; Caspari, 2022). While small proteins like Venus can enter the pyrenoid, most thylakoid membranes are excluded, leading to the observed decrease in chlorophyll autofluorescence in this spot. Venus channel signal emanating from the pyrenoid is thus a useful visual guide to true chloroplast localization of the fluorescent reporter (Caspari, 2022). Note that due to cTP-C containing a Rubisco-binding motif (Meyer et al., 2020), Venus accumulation in the Rubisco microcompartment, the pyrenoid, is particularly pronounced in constructs carrying this element. When natively R-rich RCA1-cTP and CAG2-mTP sequences were subjected to systematic R→K substitutions (^c^CH_K_^c^, MH_K_^m^), their ability to target was reduced or abolished (Fig. 3E, Fig. S18-S19). These experiments demonstrate the respective functional contributions of K and R residues to antimicrobial and organelle targeting activity.

We then systematically re-examined the organelle targeting ability of the set of 13 HA-RAMP that had undergone K→R substitutions (Fig. 2 row B, Fig. S3-S15). MII_R_ now shows much improved mitochondrial targeting (Fig. 3F), with a large majority of Venus colocalizing with MitoTracker (note that the MitoTracker occasionally appears to stain the cell envelope in addition to the mitochondria, thus not all the MitoTracker signal colocalizes with Venus; cf. Fig. S2A). R2G_R_ as well as CP3_R_ show a gain of partial targeting, as evidenced by significant overlap of Venus and MitoTracker fluorescence (Fig. 3F). Thus, an exchange of K into R increases targeting by HA-RAMP constructs. This effect is not exclusive to mitochondrial targeting (Fig. 3G): Across constructs (i.e. including those containing additional modifications), the K→R switch enabled or improved mitochondrial targeting in 8 cases (Fig. 2, rows = capital letters, columns = lower case letters; gain: ABcg, CDgi, GHf; improve: ABb, CDc, GHc) and chloroplast targeting in 15 cases, including 3 cases of dual targeting (Fig. 2, gain: CDm, EFg, GHgilm, KLg; improve: EFm, GHk, KLafj; dual: IJab, KLc). Lost or decreased targeting was observed in only three cases (Fig. 2, loss: IJgj, decrease: EFh). See Fig. S24 for illustrative examples.

### TP C-termini matter for targeting

Addition of mTP-C, the cleavage-site containing C-terminal element of CAG2-mTP enabled partial mitochondrial targeting in two constructs (R2G_K_ and E1S_R_) and improved mitochondrial targeting in a further two constructs (R2G_R_ and CP3_R_), in addition to continued targeting by B2I and MII (Fig. 2 rows CD). As an indicative example, gain of targeting in R2G_K_ is shown in Fig. 4A. The impact of adding cTP-C, the C-terminal element derived from RCA1-cTP, is even more important: cTP-C significantly enabled chloroplast targeting, which could be seen in nine HA-RAMP constructs involving CP3_R_, S1D, LCA, B15 and EHF (Fig. 2 rows E,F). Note that the addition of cTP-C was also compatible with mitochondrial localization by B2I and MII (Fig. 2 rows E,F). Gain of partial chloroplast targeting in LCA_K_ and full targeting in EHF_R_ are shown as examples in Fig. 4A. The addition of mTP-C also enabled chloroplast localization by EHF_R_ (Fig. 4A, Fig. 2 row D). Low Venus accumulation in EHF_R_ and EHF_R_^m^ means brightness needed to be set low enough that autofluorescence accounts for at least some of the signal that is colocalized with the chlorophyll channel (Fig. 4A). EHF_R_ shows Venus signal in the cytoplasm, but not within the pyrenoid, which was interpreted as an absence of targeting. By contrast, Venus signal emanating from within the pyrenoid in EHF_R_^m^ provides unambiguous evidence for chloroplast localization.

In TP controls, deletion of mTP-C reduces but does not totally abolish mitochondrial targeting of CAG2-mTP (MH_R_), while replacing mTP-C with cTP-C (MH_R_^c^) partially retargets the construct to the chloroplast (Fig. 4B; Fig. 2 column q, rows BDF). By contrast, deletion of cTP-C abolishes chloroplast targeting by RCA1-cTP (^c^CH_R_), as does replacing cTP-C with mTP-C (^c^CH_R_^m^) (Fig. 4B; Fig. 2 column p, rows HIL).

### cTP N-termini matter for chloroplast targeting

The sole addition of cTP-N generated at least partial chloroplast targeting in 6 HA-RAMP constructs, notably by SIM (Fig. 4C), but also CP3_R_, E1S_R_, B15_R_ and EHF_R_ (Fig. 2 rows G,H). However, cTP-N also enabled mitochondrial targeting in three HA-RAMPs that had not previously shown mitochondrial localization, B1E (Fig. 4C), DDM_R_ and S1D (Fig. 2 rows G,H).

The importance of cTP-N as a chloroplast determinant becomes more evident when combined with a C-terminal element, as shown in Fig. 4D using the example of ^c^B2I_R_: while ^c^B2I_R_ (partially) targets the mitochondria, addition of mTP-C (^c^B2I_R_^m^) results in dual targeting to the chloroplast in addition to the mitochondria, and addition of cTP-C (^c^B2I_R_^c^) results in targeting only to the chloroplast. Across HA-RAMP constructs, combining cTP-N with mTP-C (Fig. 2 rows I,J) resulted in 6 cases of dual targeting (B2I_R_, MII_R_, B1E, S1D) and 10 cases of chloroplast targeting (CP3_K_, E1S, LCA_K_, SIM, B15, EHF). Combining cTP-N with cTP-C (Fig. 2 rows KL) generated at least partial chloroplast targeting in 22 out of 26 HA-RAMP constructs (including 1 instance of dual targeting by R2G_R_); only DS4 fails to show any chloroplast targeting.

Finally, Fig. 4E (also Fig. 2 column q, rows JL) shows that while addition of cTP-N to the CAG2-mTP abolished mitochondrial targeting (^c^MH_R_^m^), the cTP-N/cTP-C combination (^c^MH_R_^c^) retargets to the chloroplast (see Fig. 4B for controls without cTP-N). Replacing the native amphipathic helix of RCA1-cTP between cTP-N and cTP-C with a poly-Alanine peptide of equal length (^c^AA^c^), or fusing cTP-N and cTP-C directly with no intervening peptide (^c^-^c^) both lead to partial chloroplast targeting (Fig. 4E, Fig 2 rows KL, columns rs). The two latter experiments demonstrate that there are enough determinants for recognition of the chloroplast translocon dispersed between the Nter and Cter of a cTP, to target Venus to the chloroplast, albeit with a lower efficiency than when an amphipathic helix is present in between. That the nature of the intervening peptide matters can be further seen in constructs that fail to target the chloroplast in the presence of cTP-N and cTP-C, such as RP2 (Fig. 4E), but also RP1, CP3_K_, DS4 and R2G_K_ (Fig. 2 rows KL).

### HA-RAMPs dominate targeting specificity

Considering Fig. 2 by columns reveals that organelle specificity is, to a large extent, determined by HA-RAMPs. Only 3 HA-RAMPs (B2I, R2G, MII; referred to hereafter as the mt-set) account for >70% of all constructs where mitochondrial targeting is seen. Similarly, 5 HA-RAMPs (E1S, LCA, SIM, B15, EHF; referred to as the cp-set) account for the majority (~57%) of all chloroplast targeting, and for two thirds when excluding the cTP-N/cTP-C combination that generates chloroplast targeting across most HA-RAMPs. In some instances, a set of peptide modifications may switch targeting from the mitochondria to the chloroplast, yet the major effect of modifications, i.e. exchanging K→R (Fig. 3, Fig. S24) or adding TP elements (Fig. 4), is to enhance the targeting ability to an organelle determined by the HA-RAMP primary sequence properties (Fig. 2).

### Probing cleavage of HA-RAMP-driven reporter constructs by immunoblotting

To characterize the maturation of HA-RAMP–targeted proteins upon organellar import, we performed immunoblotting experiments using whole-cell extracts probed with a FLAG antibody targeting the Venus-FLAG reporter (Fig. 5). Indicative examples were selected for clarity; a more comprehensive overview is provided in Fig. S28.

**Fig. 5.**
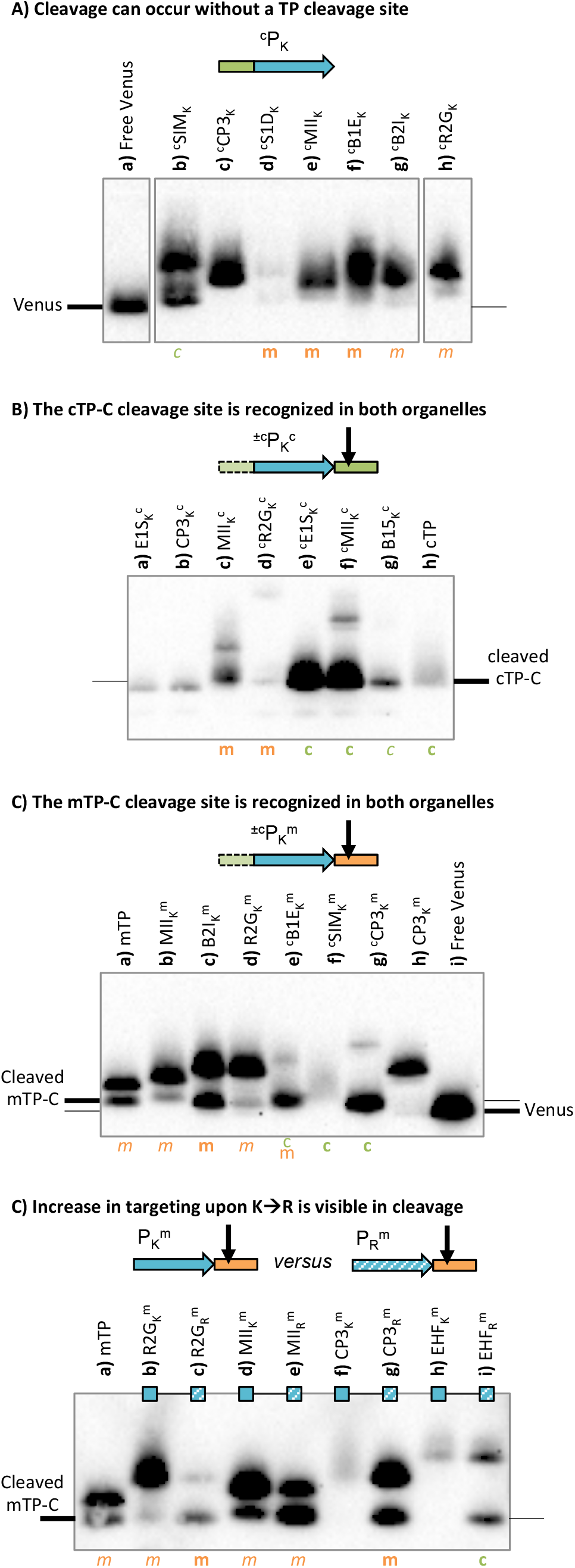
Import is associated with maturation of the preprotein. Western Blots used an α-FLAG antibody on selected constructs as indicated above blots. Where a construct was interpreted as generating reporter localization by fluorescence microscopy (cf. Fig. 2, Fig. S3-S20) in mitochondria or chloroplast, this is indicated by an orange ‘m’ or a green ‘c’ respectively, in bold for full targeting or in italics for partial targeting. In (**A**) some lanes were spliced for clarity; the uncropped blot is provided in Fig. S28. The migration of Venus without any presequence, and Venus with additional amino acids at the N-terminus left over after cleavage of cTP-C and mTP-C, are indicated for reference based on free Venus and cTP/mTP controls respectively. The cTP control is RCA1-cTP (construct ^c^CH_R_^c^), the mTP control is CAG2-mTP (construct MH_R_^m^).

In the absence of a dedicated cleavage site (Fig. 5A), we found some processing did occur but preproteins are also maintained, as evidenced by two bands being present in a given lane. The upper band corresponds to unprocessed Venus, which is a fusion of the reporter with the HA-RAMP construct and migrates at varying positions depending on the length of the presequence. The lower band corresponds to the processed form migrating closer to the “free” Venus position (lane a) depending on the exact site of cleavage. In HA-RAMP constructs, the majority of Venus remained in the unprocessed top band for constructs targeting the mitochondria (lanes d-h). Nonetheless, the presence of a faint processed form as lower bands just above free Venus shows that some processing occurs. Such processing is suggestive of import but not conclusive, since proteolysis can occur unlinked to import. The proportion of processed versus unprocessed Venus preprotein is much higher in partially chloroplast-targeted ^c^SIM_K_ (lane d), hinting at more efficient degradation of the unprocessed form in the chloroplast.

Addition of a cleavage site improves processing across organelles. In the presence of cTP-C (Fig. 5B), which contains the RCA1-cTP cleavage site, HA-RAMP constructs that show evidence of organellar targeting appear to be processed at a site corresponding to the cTP control (lane h), independently of whether the mitochondria or the chloroplast are targeted (lanes c-g). Constructs that do not target (lanes a,b; see also Fig. S28C lanes a,e,i) also appear to be processed, but at a site different to the one used in organelles, suggesting cTP-C may be recognized by cytoplasmic peptidases. Fig. 5C shows constructs equipped with mTP-C, which contains the CAG2-mTP cleavage site. Here, the mTP control (lane a) shows two bands, consistent with partial targeting observed by microscopy (cf. Fig. 3E, Fig. 4B). The top band thus likely corresponds to the preprotein and the bottom band to the mature form within the mitochondria. HA-RAMP constructs equipped with mTP-C that target either mitochondria or the chloroplast (lanes b-g) show lower bands that migrate at or near the cleaved mTP control. These results indicate that mTP and cTP cleavage sites are recognized in both organelles.

A switch from K to R (Fig. 5D), which improved targeting, also increased the amount of processed form relative to that of the unprocessed form in all cases, whether due to a gain of targeting (lanes f-i) or an increase in efficiency (lanes b-e), suggesting that cleavage can serve as a proxy for targeting. Indeed, constructs that show targeting, as judged from microscopy, also show evidence of cleavage in immunoblots: second bands are present for constructs lacking cleavage sites (Fig. S28A,D), while cleavage site-containing constructs migrate at the size expected for processed cTP-C (Fig. S28C,F) or mTP-C (Fig. S28B,E). We note that faint processed bands can be seen for several additional constructs (Fig. S28), notably among unmodified, HA-RAMP (Fig. S28A) suggesting partial targeting may occur in these cases, backed up by high image quantification values e.g. for CP3_K_ and R2G_K_ (Fig. S2EF, S5A, S9A). Such very partial targeting is however below the detection limit of our targeting assessment based on fluorescence microscopy.

### Chloroplast targeting involves longer, less helical peptides

To understand what differentiates mt-set from cp-set HA-RAMPs, we examined the sequence characteristics of our 13 HA-RAMPs with those of well-characterized Chlamydomonas targeting peptides (Fig. 6). In Chlamydomonas, cTPs are on average 49 residues in length (Fig. 6A), and significantly longer than mTPs (t-test, p=0.0017) that are on average 37 residues long. The difference is even greater than shown here given that many cTPs require a contribution from post-cleavage site residues for successful targeting (Bionda et al., 2010; Caspari, 2022). Consistently, cp-set HA-RAMPs (in green, Fig. 6A) are longer than mt-set HA-RAMPs (in orange; p=0.0184) and require further elongation by addition of TP elements before targeting can be observed (Fig. 2).

**Fig. 6.**
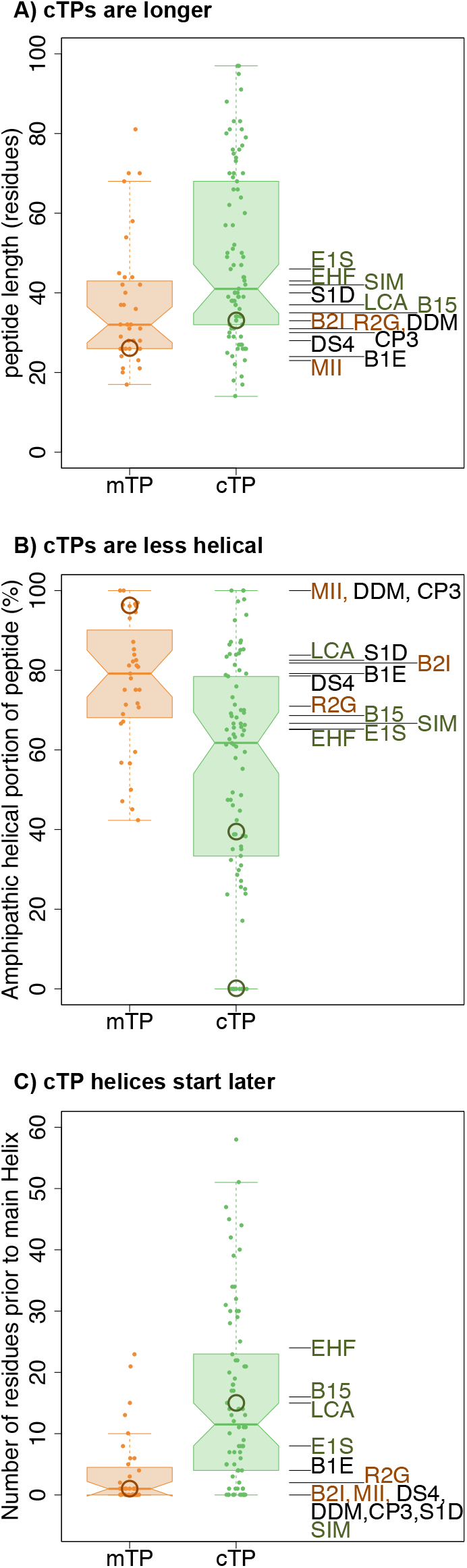
Chloroplast and mitochondrial targeting HA-RAMPs match cTPs and mTPs respectively. For salient properties, Chlamydomonas mTPs and cTPs are compared to our 13 HA-RAMPs. TP distributions are shown as boxplots (centre line: median; box limits: upper and lower quartiles; whiskers: min/max values within 1.5x interquartile range), coloured points represent individual peptides. The position of the CAG2-mTP and RCA1-cTP is circled in each graph. The non-zero value for RCA1-cTP in (B) and the single circle in (C) report on the amphipathic helix established by an NMR-study (Krimm et al., 1999), since no helix could be predicted by our approach. HA-RAMPs are colour-coded by preferred targeting: orange = mt-set; green = cp-set (cf. Fig. 2). (**A**) cTPs are significantly longer than mTPs (p=0.0017), and cp-set are longer than mt-set HA-RAMPs (p=0.0184). (**B**) Helices make up a significantly smaller fraction of cTPs than mTPs (p<0.0001), and HA-RAMPs show a similar trend (0.1122). (**C**) cTPs contain significantly longer sequence stretches upstream of the longest predicted helix than mTPs (p<0.0001), as do cp-set compared to mt-set HA-RAMPs (p=0.0205). Reported p-values were obtained through two-way t-tests for TPs and one-way t-tests for HA-RAMPs.

Most mTPs fold into an amphipathic helix for ~80% of the sequence on average, Fig. 6B) starting directly from their N-terminus (Fig. 6C). The fraction dedicated to amphipathic helix formation in cTPs is significantly lower (Fig. 6B, p<0.0001), with a longer stretch of non-helical sequence at the N-terminus (Fig. 6B, p<0.0001). In line with this contrast, 4 out of 5 of the cp-set HA-RAMPs have longer sequence stretches upstream of the main helix than the mt-set HA-RAMPs (Fig. 1, Fig. 6C). (Franzén et al., 1990) (Supplementary text, Fig. S22, Fig. S23). Also note that cTPs are predicted to be more prone to protein interaction than mTPs (Supplementary text, Fig. S26, Fig. S27).

## Discussion

### How to make TPs from HA-RAMPs

To gain insight into peptide features that govern targeting, we used a related but separate class of peptides, HA-RAMPs, as chassis and stacked modifications to generate targeting into mitochondria or chloroplasts, or in some cases into both (Fig. 7A).

**Fig. 7.**
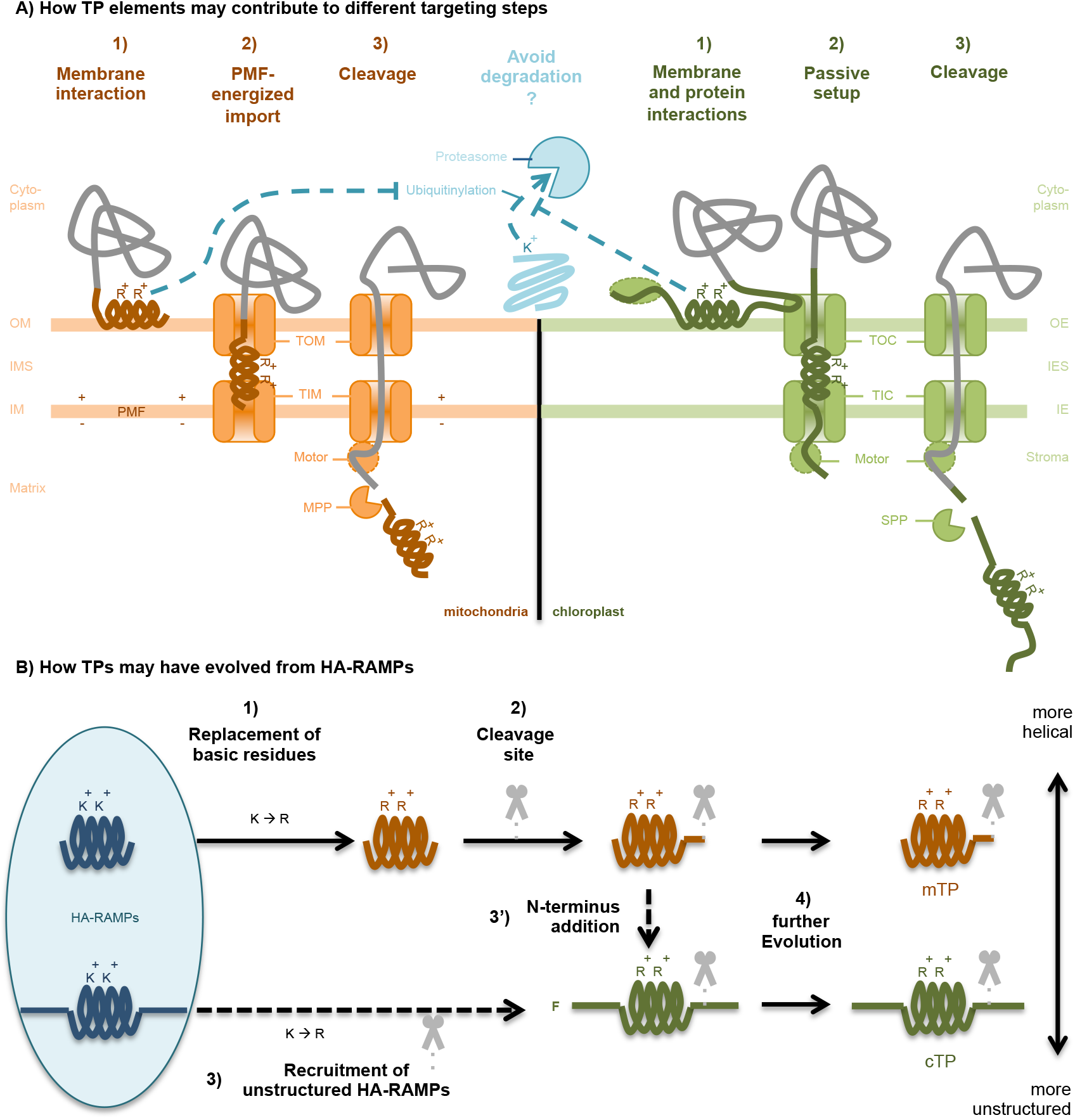
Proposed functioning and evolution of TPs. Abbreviations: OM - outer membrane, IMS - inter-membrane space, IM - inner membrane, OE - outer envelope, IES - inter-envelope space, IE - inner envelope, PMF - proton motive force, TOM - translocon of the outer mitochondrial membrane, TIM - translocon of the inner mitochondrial membrane, TOC - translocon of the outer chloroplast envelope, TIC - translocon of the inner chloroplast envelope, MPP - matrix processing peptidase, SPP - stromal processing peptidase, R - Arginine, K - Lysine, F - Phenylalanine, mTP - mitochondrial targeting peptide, cTP - chloroplast transit peptide. (**A**) A number of roles for TP elements during protein import are suggested. Since ubiquitination targets K residues, a preference for R in TPs may increase preprotein stability. R may also play a role in lipid interactions. (1) membrane interactions may play a role in enabling differential targeting due to organelle-specific lipid preferences of TP helices. Protein interactions by cTP N- and C-terminal elements, e.g. with cytosolic factors or TOC-components (subunits are not shown for simplicity), may also play a role in specific targeting. (2) Import of positively-charged mTPs across IM is energized by the PMF. By contrast, a passive setup is required for cTPs to allow N-termini to reach into the stroma and contact the motor complex, likely contributing to increased length and relatively unstructured N- and C-termini of cTPs. (3) Sequence elements contributing to targeting may be present downstream of cleavage sites in cTPs. (**B**) If TPs evolved from AMPs, 1) HA-RAMPs would have first changed K to R and 2) acquired a cleavage site to become mTPs. To generate cTPs, either 3) more unstructured HA-RAMPs were recruited directly to become cTPs by undergoing a K to R shift and acquiring a cleavage site separately, or 3’) mTPs acquired an N-terminal non-helical domain. Early cTP N-termini likely contained a starting F (Wunder et al., 2007). 4) Further evolution would have reinforced the differences between cTPs and mTPs to limit mis-targeting.

#### Arginines target better than Lysines

We showed that a K-to-R-switch increases targeting efficacy to both organelles across many constructs. R is more common than K in cTPs across green algae, vascular plants, red algae, glaucophytes and many secondary plastids (Patron and Waller, 2007), but the functional significance for targeting efficacy had not previously been recognized. Our observation is in line with a previous report that a R-to-K switch at the N-ter of an Arabidopsis mTP abolished targeting (Lee at al., 2019). The underlying mechanism warrants further research. Contributing factors may be differences in bulkiness and pKa (Li et al., 2013; Li et al., 2017), or in trans-acting factors that regulate targeting. For example, R features in consensus sequences of the cleavage sites (Tardif et al., 2012; Calvo et al., 2017); recognition of R by processing peptidases may thus potentially account for increased chloroplast-localization among cTP-N bearing constructs (Fig. 2 rows GH). Since K is used for ubiquination (Mattiroli and Sixma, 2014), a preference for R over K in TPs may help to protect preproteins from degradation while they transit in the cytosol (Fig. 7A). We note that the 20S proteasome is already present in archaea, with effectors specifically targeting K, like in the eukaryotic cytosol (Maupin-Furlow et al., 2006; Maupin-Furlow, 2013) which means that K residues were likely targets for protein degradation when primary endosymbiosis led to formation of the proto-mitochondrion.

#### Unstructured sequences contribute to chloroplast determination

Differences between cTP and mTP N-termini have previously been recognized to contribute to differential targeting (Bhushan et al., 2006), but the sequence features underlying this differentiation have been a matter of debate. In vascular plants, the presence of a N-terminal multi-Arginine motif was shown to prevent chloroplast import, leading to a proposal that this feature was solely responsible for differential targeting (Lee et al., 2019; Lee et al., 2020). A different research effort has focused on the presence of Hsp70 binding sites within cTP N-termini as crucial for enabling import (Chotewutmontri et al., 2012; Chotewutmontri and Bruce, 2015; Chotewutmontri et al., 2017). Here, we find that multiple Arginines are not uncommon in Chlamydomonas cTP N-termini (Fig. S23) and that Hsp70 binding sites are equally present in mTP N-termini (Fig. S26). We found that HA-RAMPs with intrinsically unstructured N-termini proved able to support chloroplast targeting, even in the absence of a cTP-N addition. This finding makes it extremely unlikely that differential import relies on specific peptide-receptor interactions mediated by co-evolved sequence motifs. Instead, we argue that the feature that differentiates cTPs from mTPs is the presence of an unstructured sequence upstream of the amphipathic helix in cTP N-termini. This view re-interprets the tripartite structure of cTPs (von Heijne et al., 1989), which is conserved all the way from glaucocystophyte algae to vascular plants (Köhler et al., 2015), by describing cTPs as composed of a central amphipathic helix flanked by N- and C-terminal unstructured sequence elements. This central helix has a weaker signal than the helix in mTPs (von Heijne and Nishikawa, 1991) and may only form upon membrane contact (Bruce, 2000; Garrido et al., 2020).

We also found that chloroplast targeting was further improved by unstructured C-termini. That TP C-termini are important to enable import was previously noted, with cTP and mTP C-termini thought to be functionally interchangeable (Lee et al., 2019). Here, we find that the nature of the C-terminus does matter: the more unstructured cTP-C enabled chloroplast targeting more often and more effectively than mTP-C, which contains a long predicted amphipathic helix. The need to contain unstructured sequences (von Heijne and Nishikawa, 1991) as well as the increased length of cTPs, even extending beyond the cleavage site in many cases (Bionda et al., 2010; Caspari, 2022), could be mechanistically related to the import system (Fig. 7A): whereas mitochondrial import makes use of the proton gradient to power uptake of positively charged presequences (Martin et al., 1991; Garg and Gould, 2016), energized chloroplast import requires the cTP to stretch across both TOC and TIC and contact the translocation motor (Chotewutmontri et al., 2017; Richardson et al., 2018; Nakai, 2018). Structured sequence elements can thus impede import, including the helix at the Venus N-terminus (Rekas et al., 2002). Consistent with this view, HA-RAMPs with unstructured C-termini, such as SIM, generated chloroplast targeting in the absence of an added TP-C.

Unstructured elements also likely provide protein-protein interaction motifs. For instance, TOC-interaction has been attributed to semi-conserved ‘FGLK’ motifs (Pilon et al., 1995; Lee et al., 2009; Chotewutmontri et al., 2012; Holbrook et al., 2016), although the requirement for F appears to be relaxed in Chlamydomonas (Fig. S26E) (Razzak et al., 2017). The presence of GLK sites with high predicted interactivity in cTP-N and cTP-C should contribute to the high chloroplast targeting potential of constructs equipped with both elements (Fig. S26).

#### Amphipathic helices contribute to differential targeting

While N- and C-terminal elements aid specificity determination, particularly for chloroplast import, their influence fails to explain why some HA-RAMPs target preferentially the mitochondria or exclusively the chloroplast. This observation suggests that targeting specificity also is determined by some sequence properties of the amphipathic helical elements, which in the case of HA-RAMPs serve to mediate insertion into specific target membranes. It is of note that individual mTPs and cTPs also interact with membrane bilayers (von Heijne et al., 1989; Bruce, 1998). For instance, the Rubisco small subunit cTP interacts with chloroplast-mimetic membranes only in presence of the chloroplast-specific galactolipids (Pinnaduwage and Bruce, 1996). Taken together these observations suggest that amphipathic helices may interact specifically with the membranes of the targeted organelle (Patron and Waller, 2007; Lazzaro et al., 2020). A direct interaction with the membrane bilayer, before that with proteins of the translocons, would provide a basic mechanism for a first step in differential organelle targeting (Fig. 7a). It could explain how TPs can be functionally specific whilst diverse in sequence.

### A possible series of events for the evolution of targeting peptides

A small subset of randomly chosen sequences is able to inefficiently deliver proteins into extant mitochondria, relying on amphipathic helices (Baker and Schatz, 1987; Lemire et al., 1989). Thus, the existing protein import machinery recognizes amphipathic helical peptides. However, an origin of TPs from random sequences does not explain how protein import into organelles would have spontaneously occurred in absence of the extant translocons.

Here, our use of antimicrobial peptides that harbour an amphipathic helix to further understand organelle targeting specificity, was inspired by the hypothesis that HA-RAMPs may have given rise to TPs during endosymbiotic organellogenesis because of an “import and destroy” mechanism from HA-RAMP-resistant bacteria (Wollman, 2016; Caspari and Lafontaine, 2021). In this context, combining the organelle-targeting behaviour of all those constructs that convert antimicrobial into targeting peptides can be translated into a temporal evolutionary scenario (Fig. 7B) that adds to previously proposed models (Lemire et al., 1989; Garg and Gould, 2016; Lee and Hwang, 2021). First, we found that several unaltered HA-RAMPs can deliver cargo, suggesting the evolution of mTPs from HA-RAMPs would have been straightforward. However, K→R was found to reduce toxicity and increase targeting, a dual effect that would have produced a strong selection pressure favouring this exchange as an early step. Since mitochondria appear to tolerate the presence of preproteins containing unaltered HA-RAMPs, the addition of cleavage sites that allow presequences to be degraded separately from the cargo protein (Kmiec et al., 2014) would have come as a second step.

That chloroplast targeting requires additional discriminating elements is consistent with cTPs evolving in a cell that already had mTPs. There are two possible scenarios for the origin of cTPs. Firstly, cTPs might have evolved directly from HA-RAMPs that already contained unstructured sequence elements (Fig. 7B), as seen for our cp-set HA-RAMPs. Secondly, as has been suggested earlier (Lee and Hwang, 2021), cTPs might have co-opted existing mTPs. According to the present scenario, these mTPs already contained R and a cleavage site, yet would still have been recognized by cyanobacterial HA-RAMP importers. In this case, the key innovation that generated cTPs may simply have been the addition of unstructured, possibly TOC-interacting elements to mTPs, which we showed here to be sufficient to retarget the mt-set HA-RAMPs equipped with TP C-termini at least partially to the chloroplast. The cTP N-terminus likely originally started with an F, given that in glaucophytes and rhodophytes (and many derived secondary plastids), a conserved N-terminal F plays a role in chloroplast import (Patron and Waller, 2007; Wunder et al., 2007; Köhler et al., 2015). This observation led to the idea that cTPs may have originated from a re-use of a C-terminal F-based motif involved in secretion via OMP85 beta-barrel proteins (Robert et al., 2006; Knopp et al., 2020) in (cyano)bacteria, from which TOC75 evolved. To this end, periplasmic POTRA domains responsible for substrate-recognition were proposed to have flipped orientation and now point into the host cytoplasm (Bullmann et al., 2010; Sommer et al., 2011). However, several studies of plant TOC75 have since consistently found POTRA-domains to be in the inter-membrane space, not the cytoplasm (Chen et al., 2016; Paila et al., 2016; Gross et al., 2020). Without this cytoplasmic receptor, there is no mechanism for how an N-terminal F could have acted as an import-enabling proto-cTP (Knopp et al., 2020).

Note that our proposed scenario makes no prediction about whether host/proto-organelle interactions were mutualistic or antagonistic. Either way, antimicrobial peptides are likely to have been part of the suite of tools used in these host/proto-symbiont interactions. In particular, the role of antimicrobial peptides in mutualistic symbioses (Mergaert, 2018) includes one of the best documented cases for defensive antimicrobial peptide import into bacteria (Guefrachi et al., 2015).

### Conclusion

Investigating the steps required to generate TPs from HA-RAMPs has allowed us to uncover a number of novel mechanistic insights. Notably, we discovered a role for Arginines for targeting efficacy, and delineated the contributions of N- and C-terminal elements in targeting specificity. Our work also suggests that whether due to common descent or convergence, the similarities between TPs and HA-RAMPs point to TPs interacting with membrane lipids as an early targeting step. As any evolutionary scenario, our plausible pathway from HA-RAMPs to TPs must be considered as a working hypothesis that will need to be assessed further by a series of bioinformatics and laboratory-controlled evolutionary experiments. Further understanding of peptide-lipid interactions and the phylogeny of import machinery components should shed new light on the evolution and functioning of organelle TPs.

## Materials and Methods

### Construct generation

Venus expression constructs were designed in SnapGene (v4.3.11) and generated by integrating PCR-amplified (Q5 hot start high-fidelity, #M0515, New England Biolabs) DNA fragments into plasmid pODC53 (Caspari, 2020) upstream of Venus using Gibson assembly (NEBuilder HiFi DNA assembly, #E5520S, New England Biolabs). Chlamydomonas TP sequences were amplified from genomic DNA extracted from strain T222+ (CC-5101). Templates for codon optimized HA-RAMP, RP and R→K modified TP sequences were obtained by gene synthesis (Eurofins Genomics). Correct assembly was verified by sequencing (Eurofins Genomics). Linear transformation cassettes were generated through restriction digest with *EcoRV* (New England Biolabs).

### Transformation and fluorescence screen conditions

Constructs were transformed into wild-type strain T222+ (CC-5101) using the protocol described previously (Onishi and Pringle, 2016), except using 4μl of DNA at 2μg/μl. Transformants (≥24 per construct) selected for Paromomycin resistance were grown in 200μl TAP in 96-well plates under 50μmol photons m^-2^ s^-2^ for 3-5 days and then screened for Venus expression in a fluorescence plate reader (CLARIOstar, BMG Labtech) as described previously (Caspari, 2020).

### Microscopy

Cells were grown in 96-well plates as described before (Garrido et al., 2020). Strains with suspected mitochondrial targeting were subjected to 0.1 μM MitoTracker Red CMXRos (ThermoFisher) in growth medium for 30 min in the dark and washed with TAP prior to taking images. Epifluorescence microscopy was performed with cells added into 200μl TAP medium, or SEM (250 mM Sucrose, 1 mM EDTA, in 10 mM MOPS-KOH pH 7.2) to stop swimming, in poly-l-lysine (Sigma Aldrich) coated 8-well μ-slides (Ibidi), using the following setup: microscope – Axio Observer.Z1 inverted microscope (Zeiss); objective – α Plan-Apochromat 100x/1.46 Oil DIC M27 (Zeiss); oil – immersol 518 F (Zeiss); camera – ORCA-flash4.0 digital camera (Hamamatsu); LEDs – 470 nm (chlorophyll), 505 nm (Venus) and white light (MitoTracker, filtered to >535nm using Zeiss beam splitter 423052-0104-000) in Colibri.2 LED system (Zeiss); filter cubes: filter 46HE YFP shift free (Zeiss, 520–550 nm emission) for Venus, custom-made filter set (559/34 BrightLine HC, Beamsplitter T 585 LP, 607/36 BrightLine HC; AHF Analysentechnik AG) for MitoTracker, and filter set 50 (Zeiss, 665–715 nm emission) for chlorophyll. Images were adjusted in Fiji (Schindelin et al., 2012) (ImageJ version 2.0.0) as described before (Garrido et al., 2020), and final figures assembled in PowerPoint (Microsoft PowerPoint for Mac 2011, Version 14.6.3).

### Automated Image Analysis

A custom ImageJ Macro was written to enable automated image segmentation of epifluorescence micrographs in Fiji. Briefly, in a given micrograph, fluorescence intensities were normalized and individual cells were detected using marker-controlled watershed from the MorphoLibJ library (Legland et al., 2016). For each cell, auto-thresholding the chlorophyll channel using the Huang method was used to generate a chlorophyll mask. Separately, the chlorophyll channel was subjected to a series of morphological filters and gaussian blurring followed by auto-thresholding, to finally detect round holes between 0.1-5μm within the ensuing binary image, to be saved as a pyrenoid mask. In MitoTracker images, the MitoTracker channel was subjected to morphological filtering followed by auto-thresholding with Otsu to generate a mitochondrial mask. Venus channel intensities were recorded for each compartment. For each cell, rotated images containing binary masks as extra channels were saved. Data was compiled, analysed and plotted in R, v3.6.1 (https://www.r-project.org/), using RStudio, 2022.07.2+576 (https://www.rstudio.com/).

### Western Blots

Cells were grown up in liquid culture (10ml TAP, 30μmol photons m^-2^ s^-1^, 160RPM) until late mid-log phase. 2ml Aliquots were resuspended in 30μl storage buffer (1× Roche cOmplete™ Mini proteinase inhibitor cocktail, 10 mM NaF, 0.2 M DTT, 0.2 M NaCO3) and stored at −20°C. 20μl boiling buffer (1× Roche cOmplete™ Mini proteinase inhibitor cocktail, 10mM NaF, 50 g/L SDS, 200 g/L sucrose) was added, then aliquots were boiled (50 seconds). Cell debris was removed (tabletop centrifuge, maximum speed, 15 minutes, 4°C). Chlorophyll content was estimated spectrophotometrically: 1 μg chlorophyll μl^-1^ = 0.11 × (OD_680_ – OD_770_) for 5 μl diluted in 1 ml water. Samples (10 μg chlorophyll, equal volumes) were run (overnight, room temperature, 18A) on large gels (35×27cm; resolving gel: 12% Acrylamide, 0.32% Bisacrylamide, 0.25 μl/ml Tetramethyl ethylenediamine, 250 mg/L Ammonium Persulfate, 375 mM Tris/HCl pH 8.8; stacking gel: 5% Acrylamide, 0.133% Bisacrylamide, 0.666 μl/ml Tetramethyl ethylenediamine, 666 mg/L Ammonium Persulfate, 125 mM Tris/HCl pH 6.8). Proteins (<25kDa, >75kDa) were transferred (0.1 μm nitrocellulose membranes, 1h, 0.8 A cm^-2^) as follows: cathode – 5 filter papers (FP, 3 mm, Whatman) soaked in transfer buffer 1 (40 mM Aminocaproic acid, 20% Isopropanol, 25 mM Tris/HCl pH 9.4) – gel – membrane – 2 FPs soaked in transfer buffer 2 (20% Isopropanol, 25 mM Tris/HCl pH 10.4) – 3 FPs soaked in transfer buffer 3 (20% Isopropanol, 300 mM Tris/HCl pH 10.4) – anode. Membranes were fixed (Ponceau Red), incubated (1h, room temperature) in block (30 g/L skimmed milk powder, 0.1% tween-20, 1x PBS: 140 mM NaCl, 30 mM KCl, 100 mM Na_2_HPO_4_, 15 mM KH_2_PO_4_), immunolabelled (overnight, 4°C) using α-FLAG 1° (Sigma Aldrich F1804, diluted 1:10 000 in block), washed (0.1% tween-20, 1x PBS), treated (1h, room temperature) with horseradish peroxidase-conjugated α-mouse 2° (diluted 1:10 000 in block), washed, revealed (ECL; ChemiDoc, Biorad). Blots were processed using ImageLab (Version 6.0.0 build 26, BioRad), final figures were assembled in PowerPoint (Microsoft).

### Antimicrobial activity assay

Minimum inhibitory concentration assays were performed as described before (Garrido et al., 2020).

### Sequence Data Set

TPs with experimentally-confirmed cleavage sites were recovered from proteomic studies: *Chlamydomonas reinhardtii* cTP (Terashima 2011), mTP (Tardif 2012); *Arabidopsis thaliana* cTP (Ge 2008, Rowland 2015), mTP (Huang 2009). For each peptide, we recovered the full-length protein sequence from NCBI and UNIPROT. Cytoplasmic control sequences were generated by taking N-terminal sequence stretches of random length, matching the distribution of peptide lengths observed in our Chlamydomonas TP dataset, from a random subset of Chlamydomonas proteins with a validated cytoplasmic location in UNIPROT. For PCAs and calculating amino acid frequencies, the same HA-RAMP, signal peptide and non-Chlamydomonas TP sequences were used as before (Garrido et al., 2020).

### Amphipathic helix prediction

Amphipatic α-helices are predicted as described previously (Garrido et al., 2020), following the principle of the Heliquest algorithm (Gautier et al., 2008). Briefly, the approach aims to establish the longest sequence stretch that contains identifiable hydrophilic and hydrophobic faces. The algorithm is iterated such that multiple non-overlapping helices can be found within a given peptide (Fig. 1). Consequently, helix fractions are calculated as number of residues within all predicted helices divided by the total number of residues in the peptide; for evaluating the number of upstream residues, only the longest helix was considered (Fig. 6, Fig. S25).

### ACC terms

In order to evaluate the physicochemical properties of our peptides we used the approach we previously described (Garrido et al., 2020). Briefly, each amino acid is described in terms of 3 ‘Z-scale’ values (Hellberg et al., 1987) that can be interpreted as representing a residue’s hydrophobic, steric and electronic properties. Auto-cross-covariances (ACC) between nearby residues are calculated up to a distance of four amino acids, generating a quantitative representation of a given peptide in terms of 36 ACC terms. Euclidian distances between HA-RAMP ACC term vectors and the barycentre of Chlamydomonas TPs were used as a measure of similarity (Table S1).

### Visualization

We performed principal component analysis (PCA) to visualize the relationships between peptides as described by their 36 ACC terms (Fig. S1, S22) or by their 5 salient TP properties (Fig. S25) using python package sklearn v0.22.1 (Pedregosa et al., 2011).

### Analysis of TP N-termini

TP N-termini, defined as the N-terminal 15 amino acids, were analysed (Fig. S23) as follows: Charge profiles were generated as described in the literature (Chotewutmontri et al., 2012). Hydrophobicity of TP N-termini was estimated using the Heliquest standalone application (Gautier et al., 2008). To evaluate disorder, we used IUpred2A, a software that calculates the probability for each residue of being in a disordered region (Erdős and Dosztányi, 2020). We used the default ‘Long’ setting, which has been reported to be more accurate than the alternative ‘Short’ setting (Nielsen and Mulder, 2019). The disorder of a given sequence was taken as the mean of the probability values for each residue (average residue disorder probability).

### Statistical prediction

To evaluate the predictive power of ACC terms (Garrido et al., 2020) obtained for 15 residue-long TP N-termini with regards to localization (Fig. S23) we used a binomial logistic regression classifier. We performed one hundred 5-fold cross validation runs. In each set of one hundred runs, we randomly selected sequences such that the same number of mTPs and cTPs were used. For C. reinhardtii we used 33 mTP and 33 cTP and for A. thaliana we use 29 mTP and 29 cTP sequences. We used an elastic net penalty in a saga solver with an l1-ratio (a-parameter) of 0, which is equivalent to using only ridge penalty, where all features take part in the model, and a C parameter (1/λ) of 0.1. The a and 1/λ parameters were optimised with 10-fold cross validation. Firstly, when 1/λ = 1, the best accuracy (0.82) was obtained with a between 0 and 0.09. Secondly, with a = 0, the best accuracy (0.82) was obtained with 1/λ = 0.1. A Logistic Regression model with elastic net penalty ratio of 0.15 (scikit-learn Python package v0.22.1) trained on class I HA-RAMPs (Garrido et al., 2020) and Chlamydomonas TPs was used to evaluate how similar potential HA-RAMP candidates are to TPs (Table S1). Custom scripts were written in python (v3.7.6).

### Interaction site prediction

Values for the Boman index, a quantitative proxy for whether a peptide is more likely to interact with proteins (high values) or lipids (low values), were calculated as described in the literature (Boman, 2003). Anchor2 interactivity values, a second proxy for protein interaction potential developed for disordered sequences (Mészáros et al., 2009), were calculated using the IUPred2A standalone application and webserver (Mészáros et al., 2018; Erdős and Dosztányi, 2020). Interaction sites for Hsp70 were predicted as described by Ivey and co-workers (Ivey et al., 2000) based on experimental affinity values for individual amino acids. Putatively TOC-interacting ‘FGLK’ sites were established by searching for presence of F and [P or G] and [K or R] and [A or L or V], and absence of [D and E], within each 8-residue window of a sequence, corresponding to rule 22 by Chotewutmontri and co-workers (Chotewutmontri et al., 2012) that was recommended by the authors in personal communication. ‘FGLK-1’ sites were established the same way, but requiring the presence of only three of the four positive determinants. Custom scripts were implemented in R using RStudio.

### Statistical analysis

Chlamydomonas TP distributions (Fig. 6, Fig. S26) were compared using two-sided t-tests (*n*=34 for mTPs and *n*=85 for cTPs), and associated mt-set and cp-set HA-RAMPs were compared using one-sided t-tests based on the trends set by TPs (*n*=3 for mt-set and *n*=5 for cp-set HA-RAMPs) in R using RStudio. Multiple Kruskal statistical tests are performed (same Chlamydomonas mTPs and cTPs as above, plus *n=*382 HA-RAMPs) to evaluate the distribution of amino acids (Fig. 3c, Fig. S21) in the different groups, followed by Dunn post-hoc tests (scipy v1.4.1).

## Acknowledgements

We are very grateful to Florian Schober and Katherine Madden for their effort and enthusiasm in helping us generate a quantitative image quantification dataset. Thank you to Tiffina Benhamou and Gabriel Chemin, who worked on aspects of this study with ODC during their summer internships. We also thank Prakitchai Chotewutmontri and Barry Bruce for their help in correctly replicating their Hsp70-binding and FGLK-motif finding algorithms.

The following financial support is gratefully acknowledged: the Centre National de la Recherche Scientifique and Sorbonne University for annual funding to UMR7141; the Agence National de la Recherche for (a) “ChloroMitoRAMP” ANR grant (ANR-19-CE13-0009) and (b) “LabEx Dynamo” (ANR-LABX-011), both of which provided postdoctoral support to ODC, and (c) “MATHTEST” grant (ANR-18-CE13-0027) which provided doctoral support to CG; finally, the Fondation Edmond Rothschild which provided complementary financial support to ODC and CG. The funders had no role in the design of the study; in the collection, analyses, or interpretation of data; in the writing of the manuscript, or in the decision to publish the results.

## Author contributions

FAW, IL and YC conceptualized the project and acquired funds. ODC devised, conducted and visualized the wet-lab experimental part of the investigation, and was chiefly responsible for assigning subcellular localization interpretations based on microscopic evidence. CG carried out the dry-lab/bioinformatic part of the investigation and associated visualization under the supervision of IL, writing custom software to do so (ODC contributed to the analysis of FGLK and Hsp70-binding sites). COL developed the custom imageJ macros used for automated image segmentation. ODC wrote the original draft of the manuscript. FAW, IL and ODC reviewed and edited the manuscript. All authors approved the final manuscript. The authors declare no competing interests.

## Data availability

All data are available in the main text or the supplementary materials. Custom code generated in the course of this project will be made available without restrictions upon request to the authors. All plasmids, and one independent insertion line per construct (except RP constructs), are available through the Chlamydomonas resource centre (https://www.chlamycollection.org/).

## Supporting Information

### Supplementary Text

**Fig. S1** HA-RAMP candidates cover a diversity of physico-chemical properties

**Fig. S2** Automated image analysis backs up manual targeting assessment

**Fig. S3** Biological replicates of Brevinin-2ISb

**Fig. S4** Biological replicates of Magainin 2

**Fig. S5** Biological replicates of Ranatuerin-2G

**Fig. S6** Biological replicates of Brevinin-1E

**Fig. S7** Biological replicates of Dermaseptin S4

**Fig. S8** Biological replicates of Dermadistinctin-M

**Fig. S8** Biological replicates of Cecropin-P3

**Fig. S10** Biological replicates of Sarcotoxin-1D

**Fig. S11** Biological replicates of Esculentin-1SEA

**Fig. S12** Biological replicates of Leucocin-A

**Fig. S13** Biological replicates of SI Moricin

**Fig. S14** Biological replicates of Bacillocin 1580

**Fig. S15** Biological replicates of Enterocin HF

**Fig. S16** Biological replicates of negative control Random Peptide 1

**Fig. S17** Biological replicates of negative control Random Peptide 2

**Fig. S18** Biological replicates of Rubisco activase cTP helical element control

**Fig. S19** Biological replicates of γ-carbonic anhydrase 2 mTP helical element control

**Fig. S20** Biological replicates of no-peptide and A-screen controls

**Fig. S21** Comparison of amino acid frequencies reveals K/R shift

**Fig. S22** PCAs reveal that N- but not C-termini differ between cTPs and mTPs

**Fig. S23** Algal and plant cTP N-ter share physicochemical differences against mTP

**Fig. S24** K→R generally improves HA-RAMP targeting

**Fig. S25** HA-RAMP properties determine their targeting propensities

**Fig. S26** Higher protein interactivity predicted for cTPs than mTPs

**Fig. S27** Hsp70 and (F)GLK sites are present in HA-RAMPs

**Fig. S28** Western Blots for K-bearing constructs

**Table S1** Similarity of HA-RAMPs to TPs quantified (Excel file)

**Table S2** Numeric values derived of bioinformatic analyses (Excel file)

**Table S3** Automated image quantification data (Excel file)

