## Supplementary Text and Figures for "Converting antimicrobial into targeting peptides reveals key features governing protein import into mitochondria and chloroplasts"

The following Supporting Information is available in this document:

**Supplementary Text**

**Fig. S1** HA-RAMP candidates cover a diversity of physico-chemical properties

**Fig. S2** Automated image analysis backs up manual targeting assessment

**Legend for Fig. S3-S20**

**Fig. S3** Biological replicates of Brevinin-2ISb

**Fig. S4** Biological replicates of Magainin 2

**Fig. S5** Biological replicates of Ranatuerin-2G

**Fig. S6** Biological replicates of Brevinin-1E

**Fig. S7** Biological replicates of Dermaseptin S4

**Fig. S8** Biological replicates of Dermadistinctin-M

**Fig. S8** Biological replicates of Cecropin-P3

**Fig. S10** Biological replicates of Sarcotoxin-1D

**Fig. S11** Biological replicates of Esculentin-1SEA

**Fig. S12** Biological replicates of Leucocin-A

**Fig. S13** Biological replicates of SI Moricin

**Fig. S14** Biological replicates of Bacillocin 1580

**Fig. S15** Biological replicates of Enterocin HF

**Fig. S16** Biological replicates of negative control Random Peptide 1

**Fig. S17** Biological replicates of negative control Random Peptide 2

**Fig. S18** Biological replicates of Rubisco activase cTP helical element (CH) control

**Fig. S19** Biological replicates of γ-carbonic anhydrase 2 mTP helical element (MH) control

**Fig. S20** Biological replicates of no-peptide and A-screen controls

**Fig. S21** Comparison of amino acid frequencies reveals K/R shift

**Fig. S22** PCAs reveal that N- but not C-termini differ between cTPs and mTPs

**Fig. S23** Algal and plant cTP N-ter share physicochemical differences against mTP

**Fig. S24** K→R generally improves HA-RAMP targeting

**Fig. S25** HA-RAMP properties determine their targeting propensities

**Fig. S26** Higher protein interactivity predicted for cTPs than mTPs

**Fig. S27** Hsp70 and (F)GLK sites are present in HA-RAMPs

**Fig. S28** Western Blots for K-bearing constructs

The following Supporting Information is available in a separate Excel file:

**Table S1** Similarity of HA-RAMPs to TPs quantified

**Table S2** Numeric values derived of bioinformatic analyses

**Table S3** Automated image quantification data

### Supplementary Text

#### Automated Image Analysis corroborates targeting

To assess subcellular Venus localisation independently of our manual targeting assignment, 2202 3-channel images containing Venus, chlorophyll and brightfield channels, and 557 4-channel images containing an additional MitoTracker channel, were subjected to automated, quantitative image analysis (Fig. S2). For reference, key strains with known targeting are shown in Fig. S2A. In the absence of Venus, low chlorophyll autofluorescence is visible in the Venus channel; in the absence of a presequence, Venus accumulates in the cytoplasm, targeting neither to the mitochondria nor to the chloroplast. When equipped with the chimeric $B15_K^c$ presequence, Venus accumulates in the chloroplast, while with $MII_K^c$, Venus is in the mitochondria as established previously using confocal microscopy and biochemistry on isolated organelles (Garrido et al., 2020). When targeted to the chloroplast, Venus not only colocalizes with chlorophyll but also accumulates within the pyrenoid. The pyrenoid can be detected as a region of lower chlorophyll fluorescence at the base of the cup-shaped chloroplast, as visualized in Fig. S2B where the pyrenoid is labelled fluorescently using an RBCS-Venus fusion protein. Given that chlorophyll autofluorescence renders colocalization of Venus channel signal with the chlorophyll channel potentially misleading in constructs with low Venus accumulation, we chose to focus on evaluating Venus signal within the pyrenoid for automated image segmentation (Fig. S2C inset). Rather than comparing pyrenoidal Venus to the entire cell, we opted for a comparison with Venus in the rest of the chloroplast excluding the pyrenoid (the chlorophyll region), since strong Venus channel signal from within the pyrenoid should generate high values for a pyrenoid/chlorophyll ratio when Venus is in the chloroplast and low values when Venus is elsewhere. This approach was able to separate chloroplast-targeting constructs very effectively from constructs where Venus localized elsewhere in the cell (Fig. S2C). Only a small region of overlap between the two categories remains (highlighted in blue). Among constructs targeting elsewhere, constructs found in this overlap region typically show low Venus accumulation, e.g. the no-Venus control falls also falls in this region (black diamond). While the ambiguity represented by this overlap region means that automated image segmentation is thus not quite at the level where subcellular localization can be reliably assigned automatically, the analysis provides strong support for our manual assignment being correct in most cases.

A proxy for automated analysis of mitochondrial targeting are shown in Fig. S2D. The graph is based on automated image segmentation, using the MitoTracker channel to find mitochondria and comparing the Venus channel signal in mitochondria to that across the whole cell. This approach show that the distribution of constructs that were manually assigned as mitochondrial-targeting scores significantly higher than the distribution of constructs with non-mitochondrial assignments. There is more overlap between these categories than was observed for the chloroplast-targeting analysis in Fig. S2C. Both the no-peptide control known to have cytoplasmic Venus (white diamond), and the $MII_K^c$ construct known to target the mitochondria (orange diamond) fall into this overlap region. Thus while the automated analysis provides support for the overall accuracy of our targeting assignment, manual targeting assignment is required for mitochondrial targeting to be determined. The fact that automated image analysis is noisier when it comes to mitochondrial targeting compared to chloroplast targeting likely stems from the fact the MitoTracker dye generates highly variable staining both within and between samples. Note also that the mitotracker appears to occasionally label non-mitochondrial structures, particularly on the outside of the cell, as is apparent for the $MII_K^c$ construct in Fig. S2A where Venus is known to label mitochondria.

The highest non-targeting and lowest targeting constructs, defining the overlap regions of (C) and (D), are shown in Fig. S2E, along with one dual-targeting construct present in the overlap region of both graphs.

**HA-RAMPs and TPs display a few differences in their amino-acid content.**
HA-RAMPs are richer in helix-breaking Glycines (G) but poorer in Prolines (P), whereas TPs are enriched in Serines (S) compared to HA-RAMPs (Fig. S20). Most of the other amino acid frequencies are broadly similar between the two sets of peptides and within range of the average value across all entries in the UNIPROT database (red lines).

***In silico* comparison of C and N-termini confirms the importance of the N-terminal domain for chloroplast determination**
To further explore the role of the N-terminal cTP domain, we performed some *in silico* comparisons of the studied peptides. In Principal Component Analyses on ACC Z-scales (Garrido et al., 2020) comparing TP N- and C-termini with our 13 HA-RAMPs (Fig. S21), cTPs and mTPs could be differentiated much better by N- compared to C-termini, supporting N-termini as important specificity determinants. Furthermore, even though algal cTP N-termini are more charged, less hydrophobic and more disordered than higher plant cTPs (Fig. S22) in line with algal cTPs being more mTP-like (Franzén et al., 1990), Chlamydomonas cTP N-termini are recognized as chloroplast-targeting in a model trained on Arabidopsis TP and vice versa (Fig. S22).

**Principal component analysis shows that cp-set HA-RAMPs and mt-set HA-RAMPs coherently share the same properties with cTPs and mTPS, respectively**
A PCA analysis based on these properties, plus the fraction of R and K, recapitulates the combined properties within each peptide (Fig. S24). The PCA shows that cp-set HA-RAMPs are localized with cTPs in the upper left area of the graph, consistent with a lower helix fraction (Fig. 1, Fig. 5b) due to longer sequence stretches upstream of the main helix than in the mt-set HA-RAMPs (Fig. 1, Fig. 5c, $p=0.0199$). Consistent with the importance of these helical features, LCA, the most helical among cp-set HA-RAMPs also shows the highest fraction of only partial chloroplast targeting (3 out of 5 targeting constructs, Fig. 2 column j rows E,F,K). Furthermore, SIM and E1S, the two cp-set HA-RAMPs with the shortest pre-helix segments (Fig2 row C), both require addition of cTP-N for chloroplast targeting (Fig. 2 columns i,k).

**cTPs are predicted to be more prone to protein interaction than mTPs**
cTPs appear more likely to interact with proteins than mTPs (Fig. S25), just like cp-set HA-RAMPs. 'FGLK' motifs that are reportedly TOC interaction sites in plant cTPs (Chotewutmontri et al., 2017) appear shortened to 'GLK' motifs in Chlamydomonas but they are associated with an increased protein interaction potential in cTPs compared to mTPs ($p<0.0001$), including within cTP-N and cTP-C elements. However, Hsp70-interaction sites (Ivey et al., 2000), which are commonly found at cTP N-termini (Chotewutmontri et al., 2017), also occur at a high frequency in mTP sequences and HA-RAMPs (Fig. S25, S26).

**Fig. S1. HA-RAMP candidates cover a diversity of physico-chemical properties.** A Principal component analysis (PCA) based on auto-cross-correllated (ACC) Z-scale values reflects divergent physico-chemical properties of Signal Peptides (SP: bacterial secretory peptides – bSP, thylakoid signal peptides – tSP, eukaryotic signal peptides – eSP) relative to Targeting Peptides (TP: chloroplast transit peptides – cTP, mitochondrial targeting peptides – mTP) and helical-amphipathic ribosomally-produced antimicrobial peptides (HA-RAMP, shown here are class I HA-RAMPs *sensu* (Garrido et al., 2020). Axes are principal components (PC) 1 and 2. Each dot represents one peptide; the 13 HA-RAMP candidates studied in this article are highlighted in red. Convex areas include the 50% of peptides at the centre of each distribution.

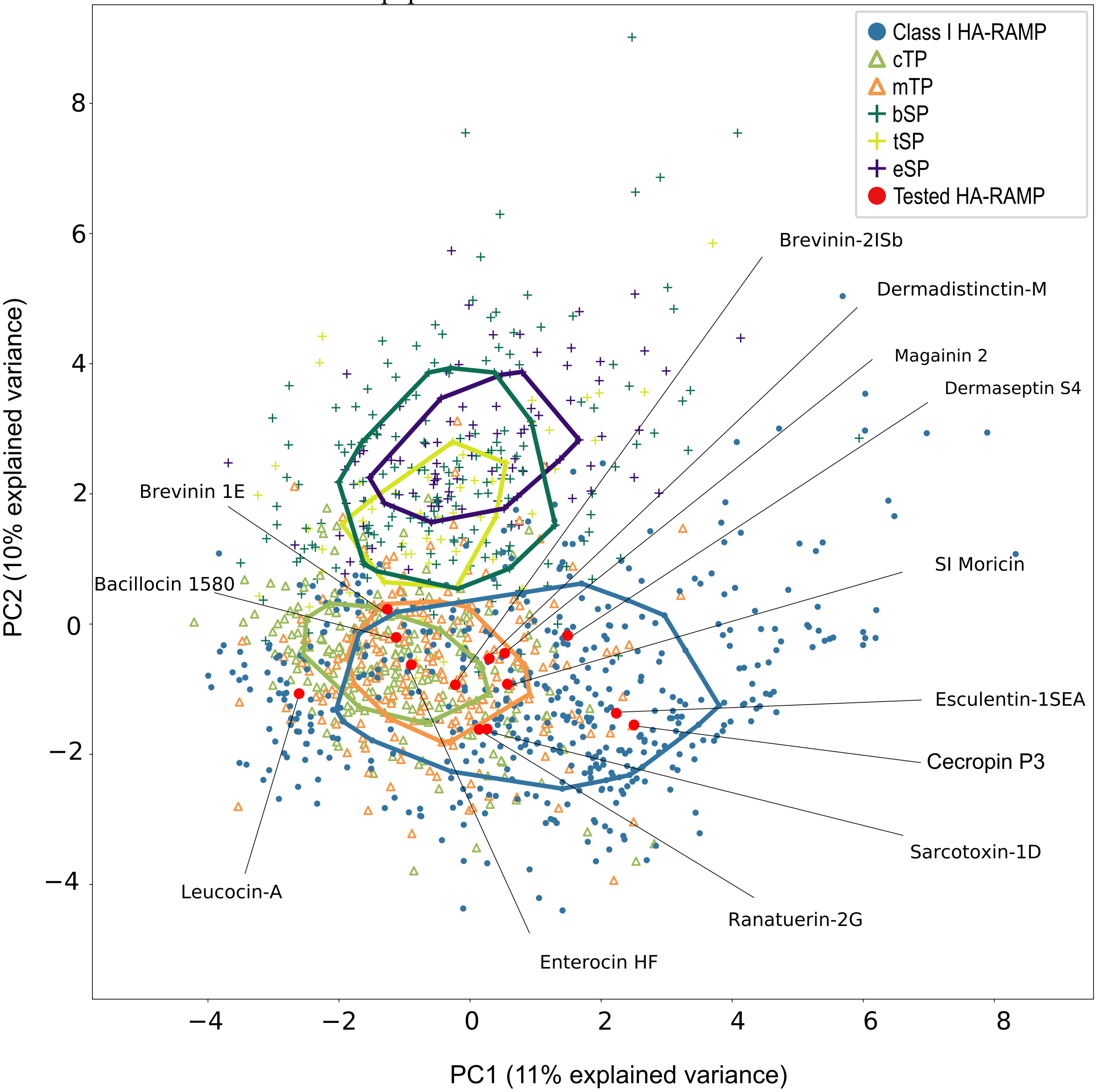

**Fig. S2. Automated image analysis backs up manual targeting assessment.** **(A)** As previously established (Garrido et al., 2020), in the absence of Venus, Venus without a pre-sequence is dispersed throughout the cytoplasm; $B15_K^c$ directs Venus import into the chloroplast; $MII_K^c$ targets the mitochondria; and low chlorophyll autofluorescence is visible in the Venus channel. **(B)** The dip in chlorophyll fluorescence at the base of the chloroplast corresponds to the pyrenoid, visualized here by labelling the pyrenoid using an RBCS-Venus fusion (Mackinder et al., 2016; Caspari, 2022). **(C)** For image quantification, Automated image segmentation on normalized fluorescence micrographs was used to define a 'chlorophyll' compartment based on the chlorophyll fluorescence channel, and a 'pyrenoid' compartment by finding holes in a separate chlorophyll mask (inset). Given that in our chloroplast-targeting constructs Venus accumulates in the pyrenoid, which is simultaneously the region with the lowest autofluorescence signal within the chloroplast, we chose to evaluate pyrenoidal Venus signal as a proxy for true chloroplast targeting. Venus channel signal in pyrenoid and chlorophyll compartments was normalized by compartment areas for each cell, and average pyrenoid/chlorophyll ratios across all cells per construct were further normalized to spread from 0 to 1. The distribution of constructs that were assigned as targeting the chloroplast is shown in green. Individual constructs are plotted within the distribution (with random y-values). This includes constructs that target the chloroplast fully (dark green) or partially (light green), as well as constructs that show dual targeting (purple). The distribution of constructs that were assigned as not targeting the chloroplast is shown in black and includes constructs targeting the mitochondria fully (dark orange) or partially (light orange), as well as constructs that target neither the chloroplast nor the mitochondria (white); the 'No Venus control' is shown as black diamond. **(D)** The graph shows the results of automated segmentation, used to define a 'mitochondria' compartment based on the MitoTracker channel, and a 'cell' compartment based on Venus and chlorophyll channels (inset). The analysis is less discriminatory than that for chloroplast targeting, likely because use of the MitoTracker dye is less reliable than expression of fluorescent proteins and results in large variation in label quality both within and between samples. Normalisations detailed for (C) apply. The distribution of constructs that were assigned as targeting the mitochondria is shown in orange, and includes constructs showing full or partial mitochondrial targeting and dual targeting. The distribution of constructs that were assigned as not-mitochondrial are shown in black and include fully/partially chloroplast targeting constructs and constructs targeting neither. In both **(C)** and **(D)**, constructs with micrographs shown in other panels of this figure are indicated with special point shapes: diamond - (A), square, triangle - (E). The statistical difference between distribution means is given (student's t-test). The region of overlap that contains both targeting and non-targeting constructs is highlighted in blue below the graph. **(E)** Example images for constructs drawn from the overlap regions in (C, D) demonstrate that targeting can be reliably assigned by visual inspection, even when automated image quantification is ambiguous.

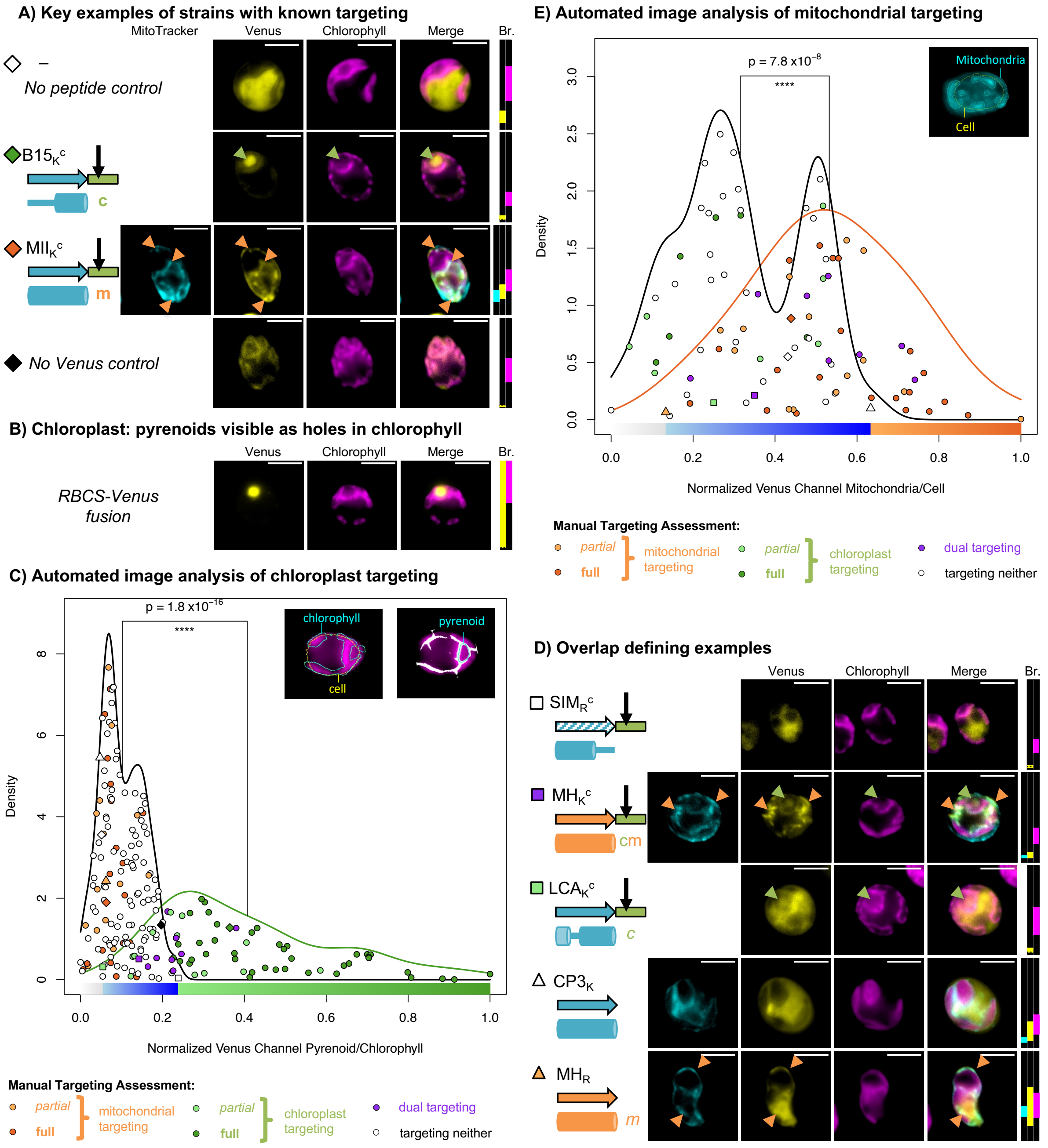
A) Key examples of strains with known targeting
MitoTracker
Venus
Chlorophyll
Merge
Br.
–
No peptide control
B15K c
c
MIIK c
m
No Venus control
B) Chloroplast: pyrenoids visible as holes in chlorophyll
RBCS-Venus fusion
C) Automated image analysis of chloroplast targeting
p = 1.8 x10−16
****
chlorophyll
cell
pyrenoid
Density
Normalized Venus Channel Pyrenoid/Chlorophyll
Manual Targeting Assessment:
partial
full
mitochondrial targeting
chloroplast targeting
dual targeting
targeting neither
E) Automated image analysis of mitochondrial targeting
p = 7.8 x10−8
Mitochondria
Cell
Normalized Venus Channel Mitochondria/Cell
D) Overlap defining examples
SIMR c
MHK c
cm
LCAK c
CP3K
MHR

**Legend for Fig. S3 – S20.** Three independent transformant lines (**Strains 1-3**) are shown for each combination of modifications (A-L), represented by a cartoon and a shorthand description (cf. Fig. 2). Where a construct was interpreted as generating reporter localization in mitochondria or chloroplast, this is indicated by an orange 'm' or a green 'c' respectively, in bold for full targeting or in italics for partial targeting. Epifluorescence microscopy images of typical cells for each strain are shown. **MitoTracker** fluorescence, false-coloured in cyan, indicates the position of mitochondria (although parts of the cell exterior are sometimes also stained). False-coloured yellow fluorescence from the **Venus** channel reports on the subcellular localization of the fluorescent reporter. **Chlorophyll** autofluorescence, shown in magenta, indicates the location of the chloroplast. Scale bars are 5µm. **Brightness** (Br.) was adjusted for clarity: fluorescence intensity values were restricted to the range shown for each channel by matching coloured rectangles. Intensity scales to 0 at the bottom of the panel, and to 65535 at the top.
On the right-hand side, quantitative information associated with each construct is shown. Data was normalized such that averages range from 0 to 1 across all constructs (cf. Fig S2). Individual cell measurements may exceed this range; for clarity, the range shown here is limited to -0.5 to 1.5, with outliers that have more extreme values shown as lying beyond a dashed line. Strain 1 is plotted as circles, strain 2 as squares, and strain 3 as diamonds. **Venus**: *Fluorescence plate reader data* of transformants screened for this construct (Venus fluorescence, accounting for cell density using linear regression of $OD_{750}$ and chlorophyll fluorescence trained on 'no venus' control transformants). Transformants that were not selected after screening are shown as small grey circles. A colour code ranging from white (low Venus fluorescence) to black (high Venus fluorescence) is used for strains 1-3 and the average thereof. *Automated image quantification data*: **pyr/chl**: Venus channel signal in the pyrenoid relative to in the chlorophyll region; **mt/cell**: Venus channel signal in mitochondria relative to the whole cell. Each point corresponds to a single cell. The colour code code is the same as that below distributions in Fig. S2, i.e. blue highlights the overlap region that contains both targeting and non-targeting constructs.

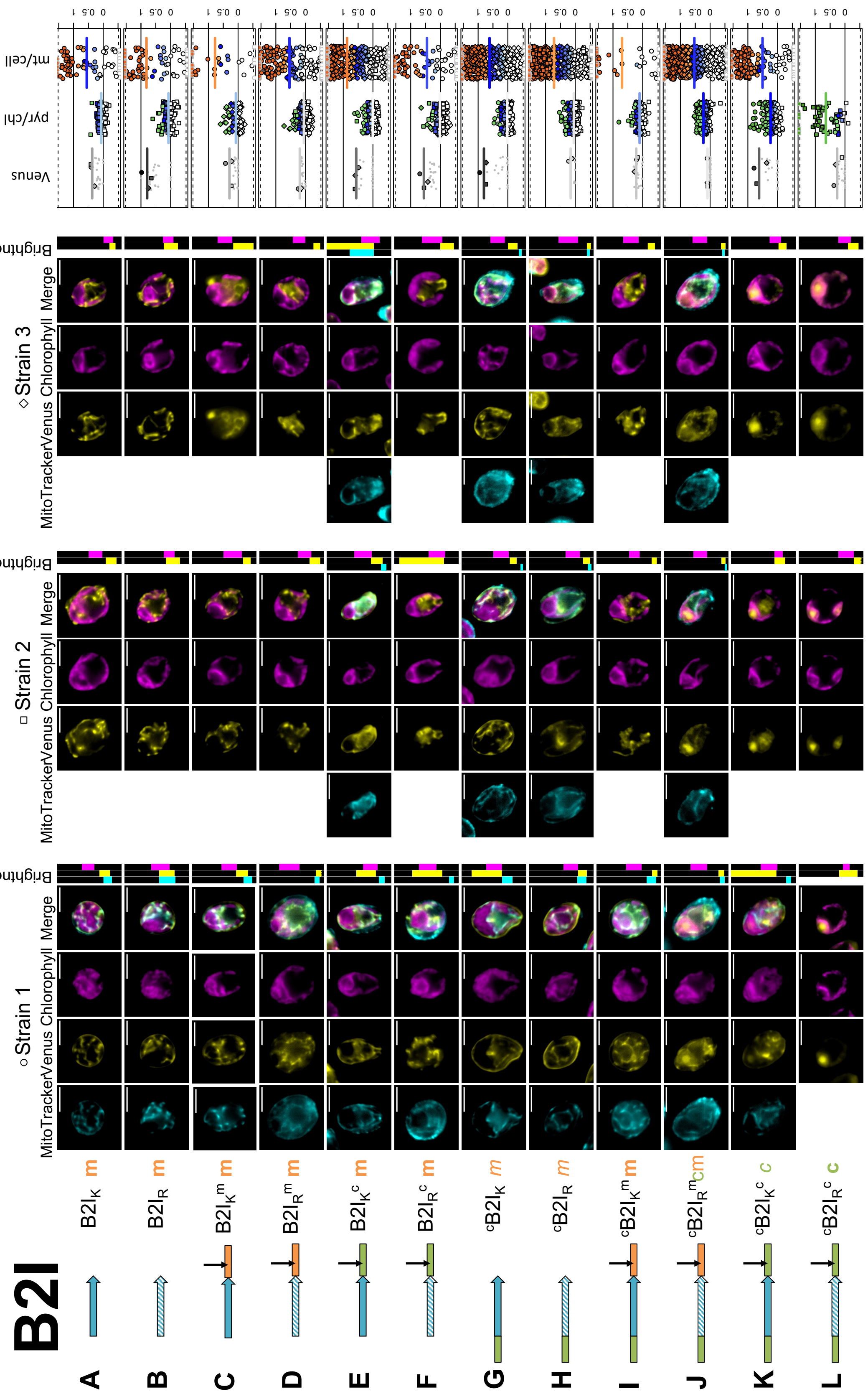


**Fig. S3. Biological replicates of Brevinin 2ISb.**

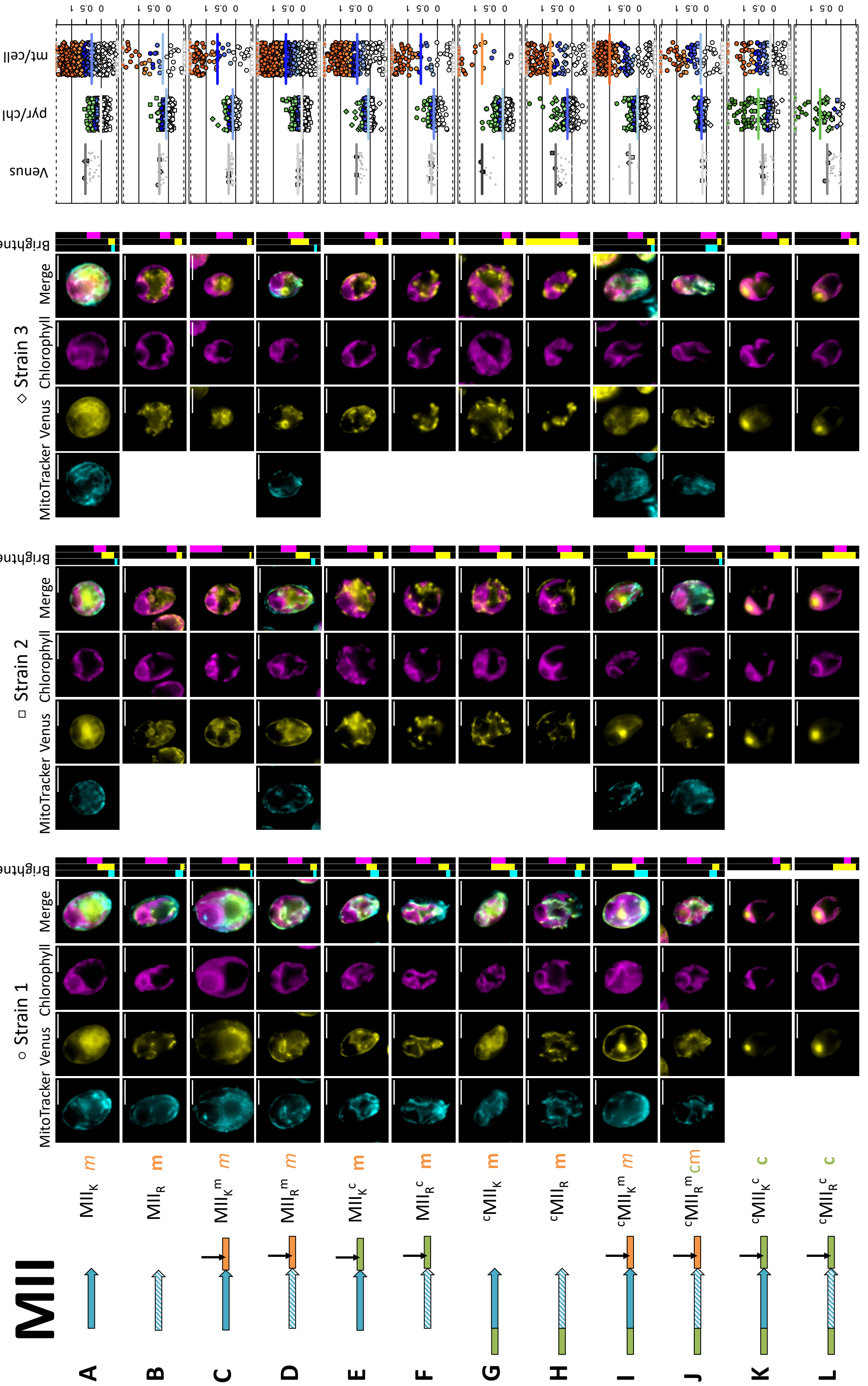


**Fig. S4. Biological replicates of Magainin 2.**

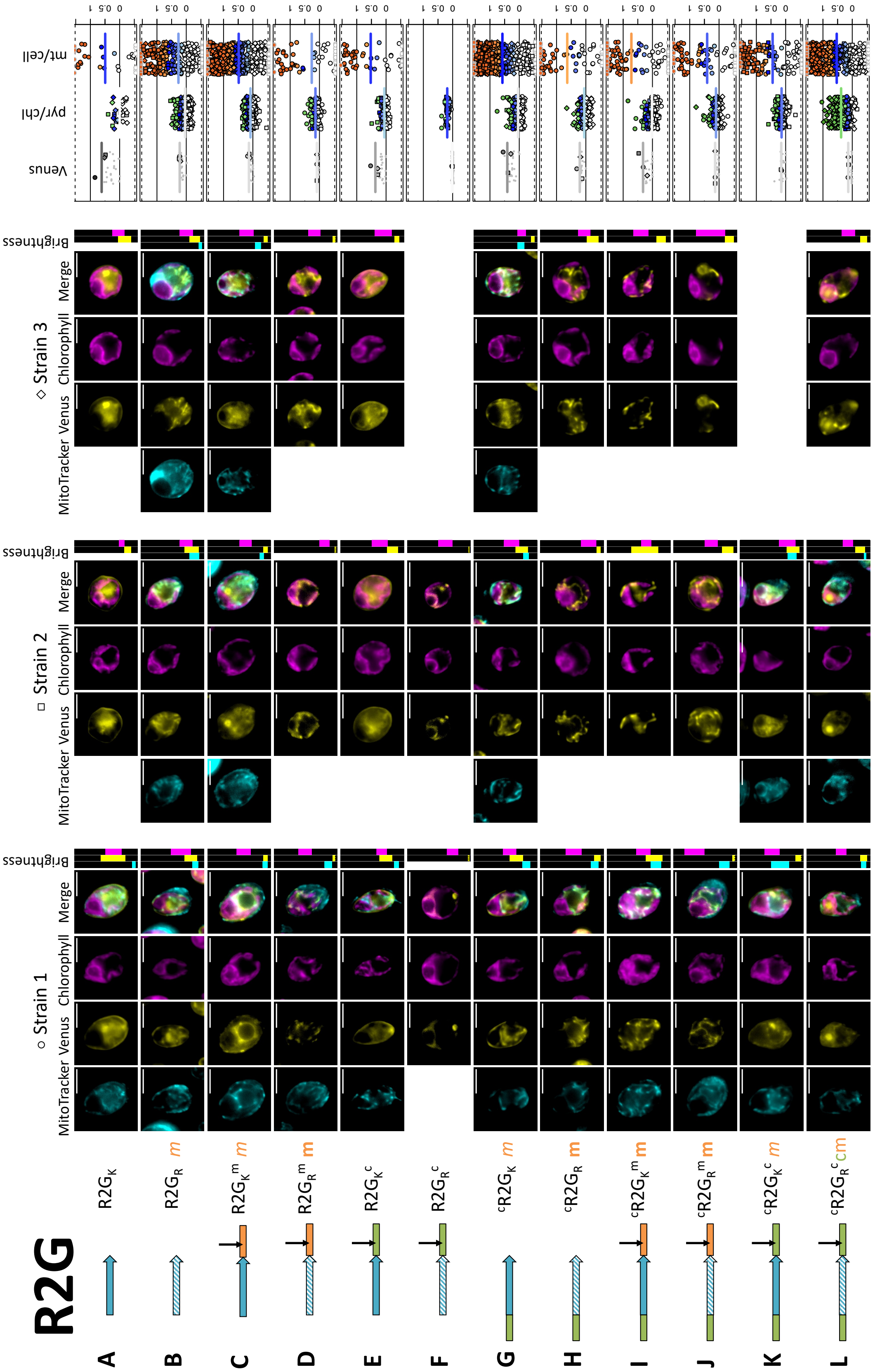


**Fig. S5. Biological replicates of Ranatuerin 2G.**

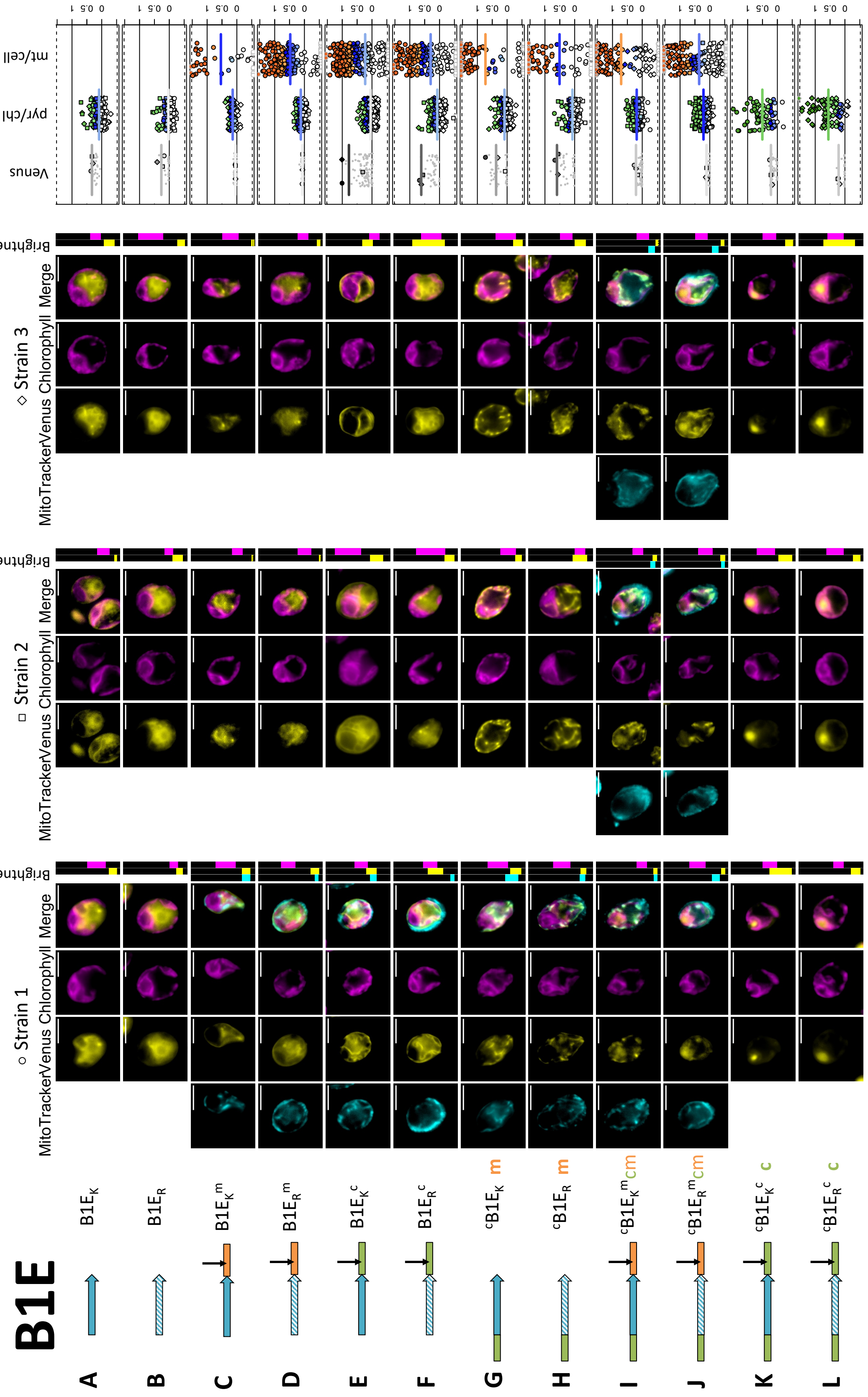


**Fig. S6. Biological replicates of Brevinin 1E.**

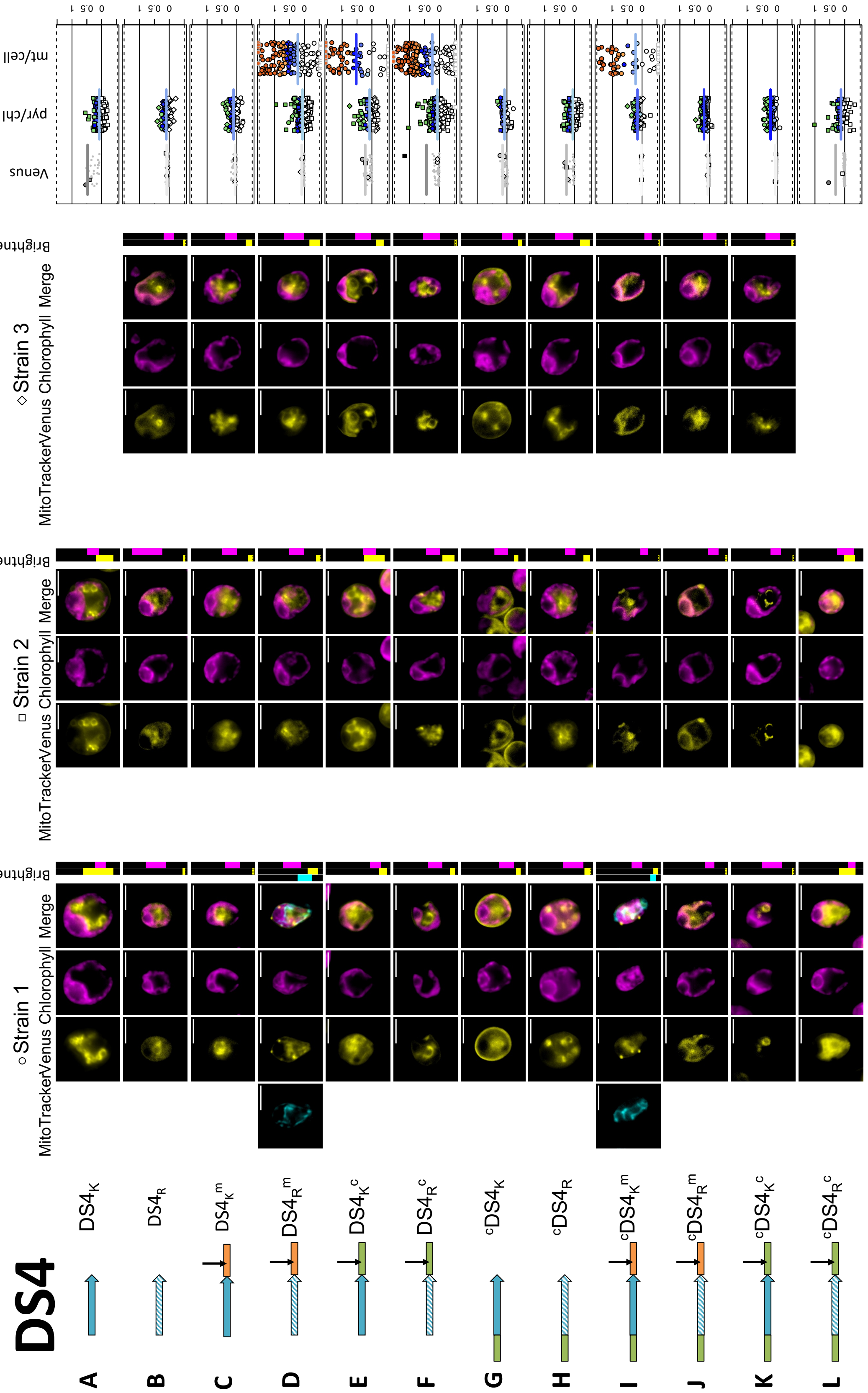


Fig. S7. Biological replicates of Dermaseptin S4.

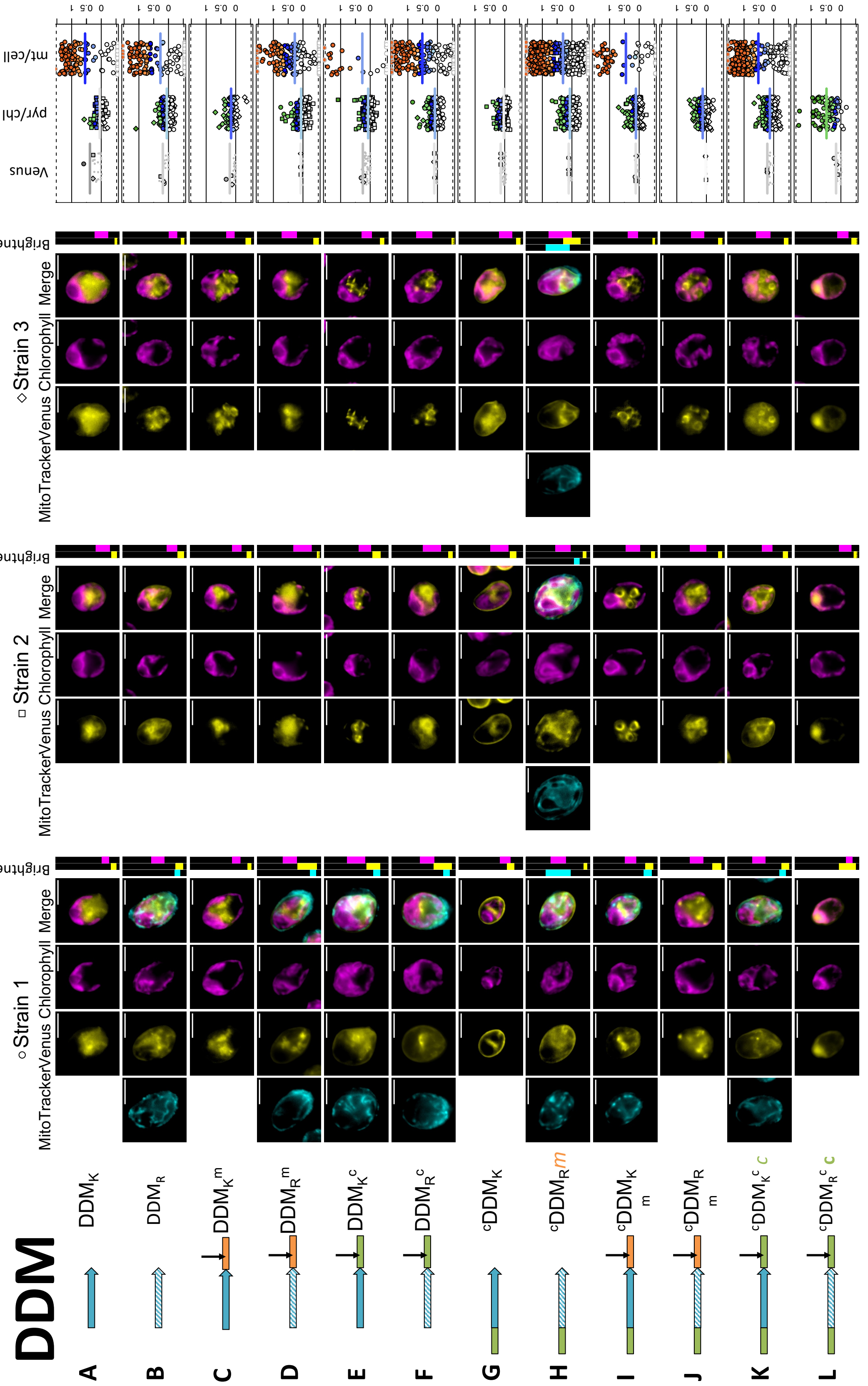


**Fig. S8. Biological replicates of Dermadistinctin M.**

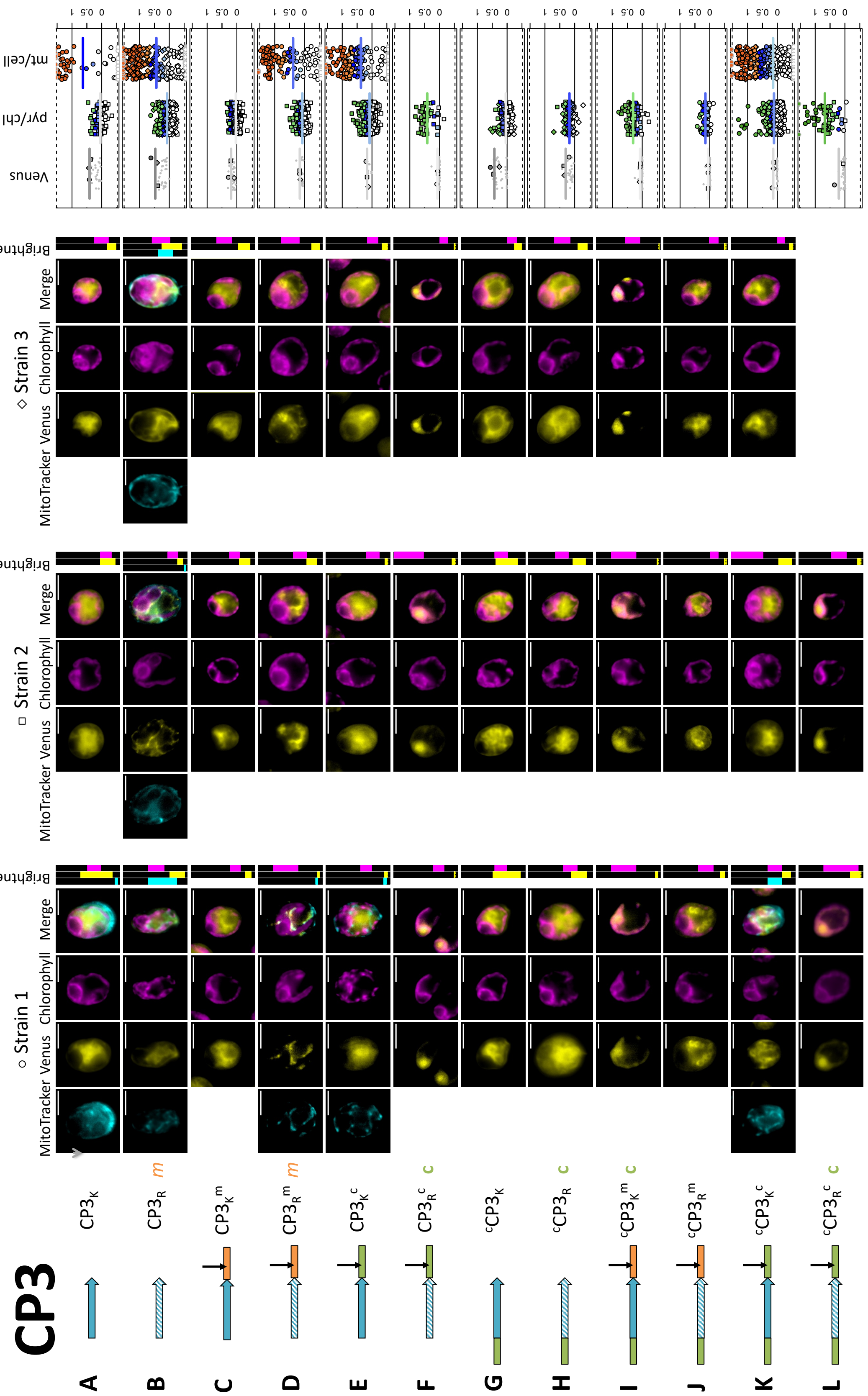


**Fig. S9. Biological replicates of Cecropin P3.**

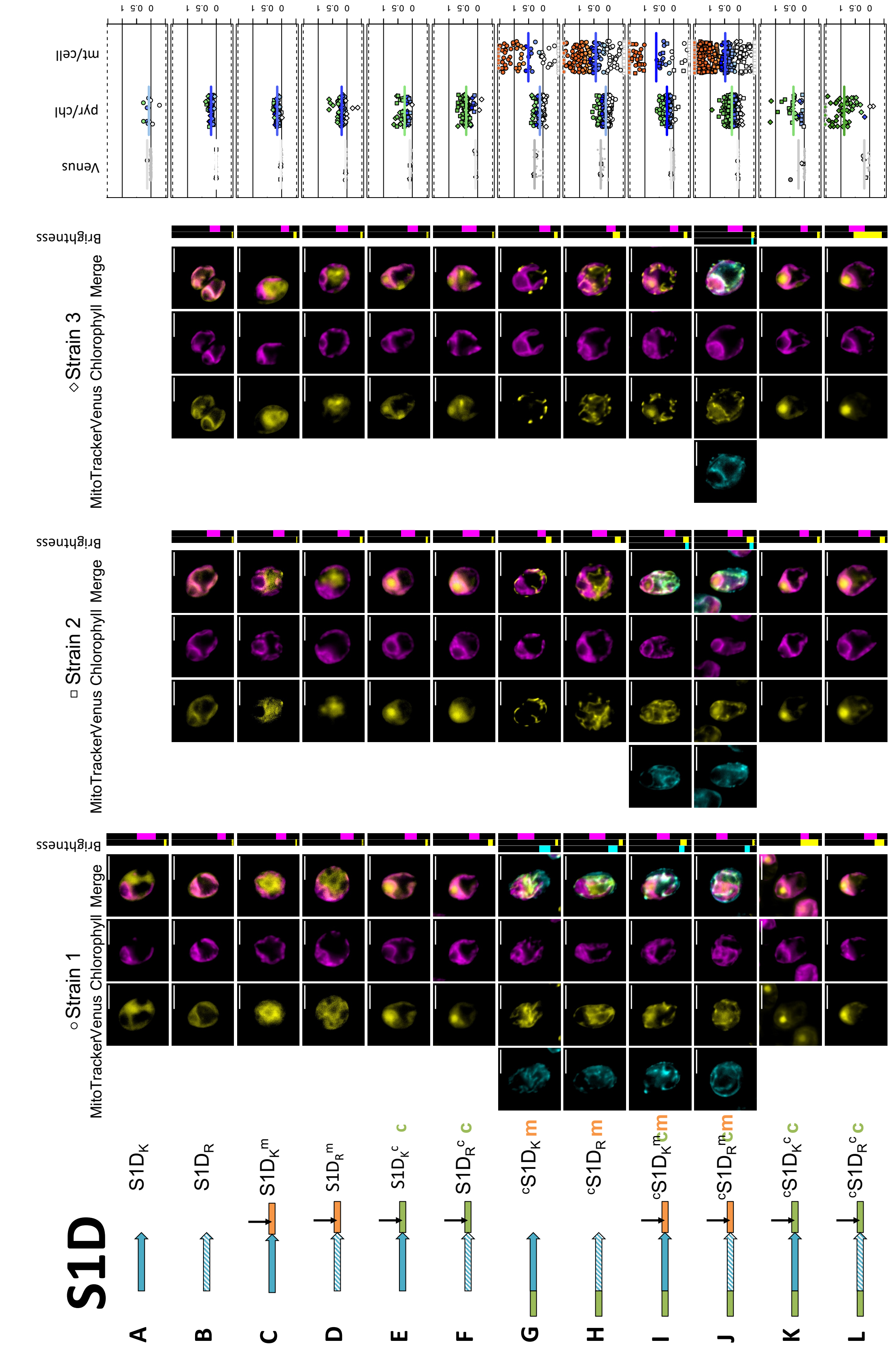


**Fig. S10. Biological replicates of Sarcotoxin 1D.**

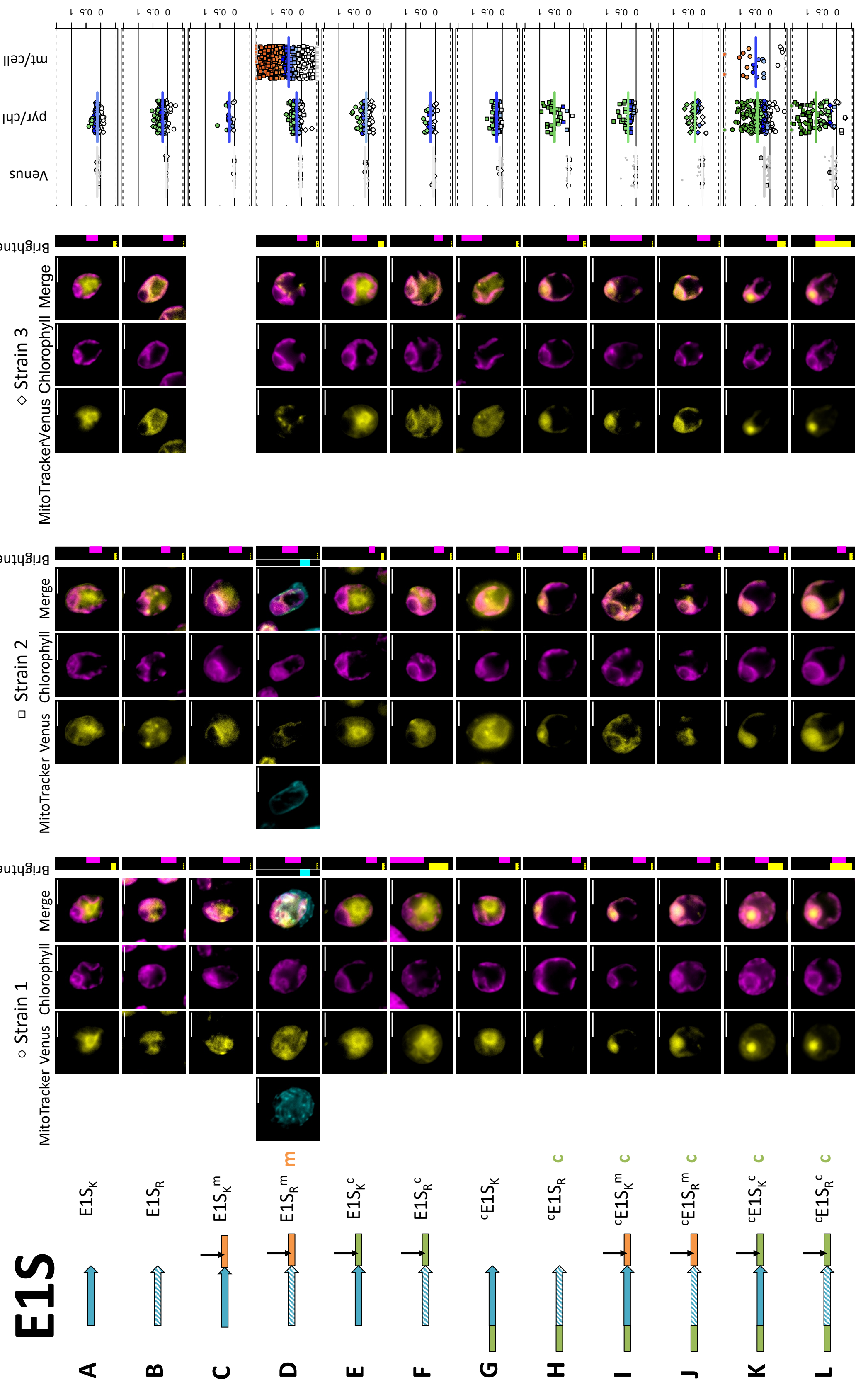


**Fig. S11. Biological replicates of Esculentin 1S.**

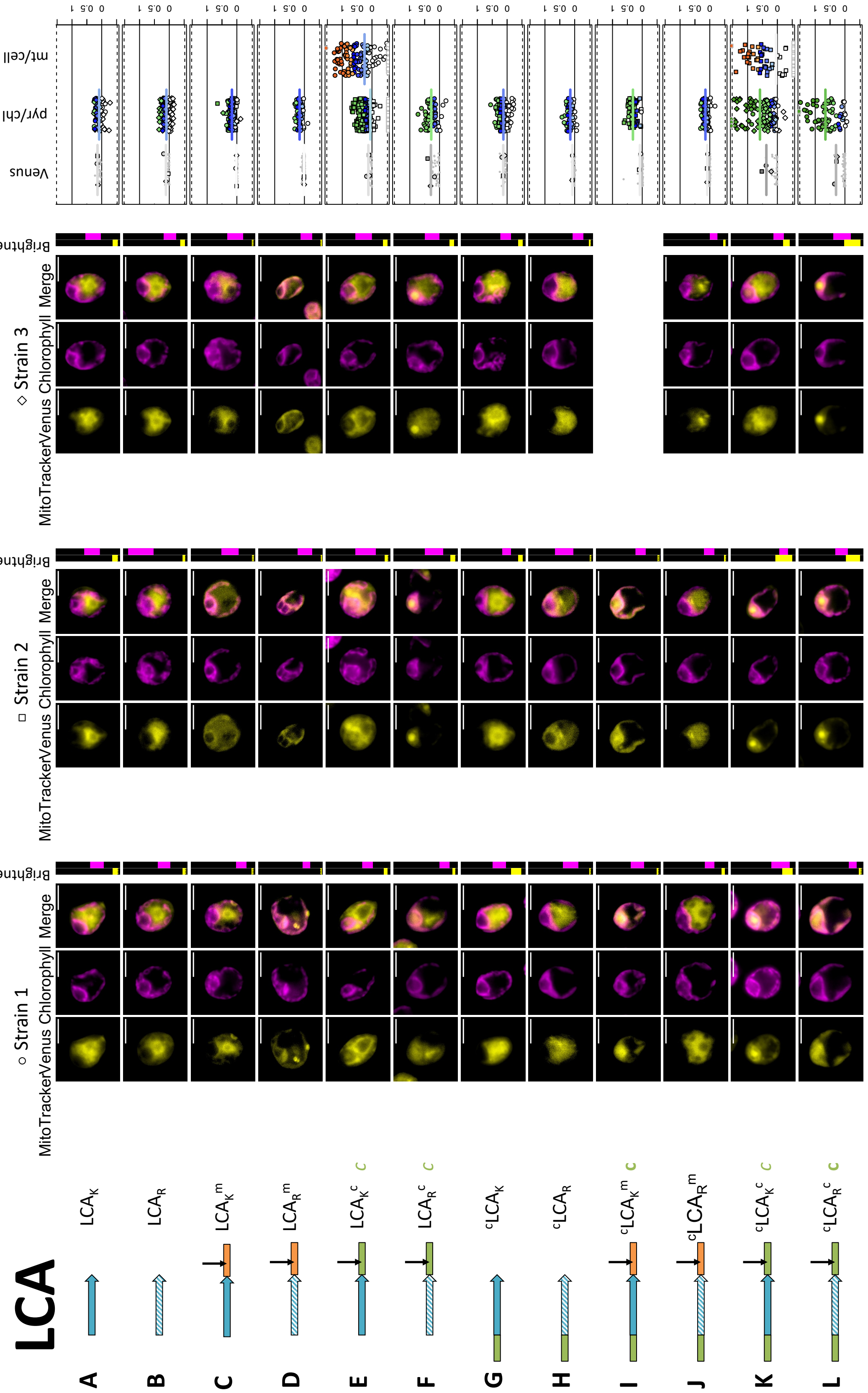


**Fig. S12. Biological replicates of Leucocin A.**

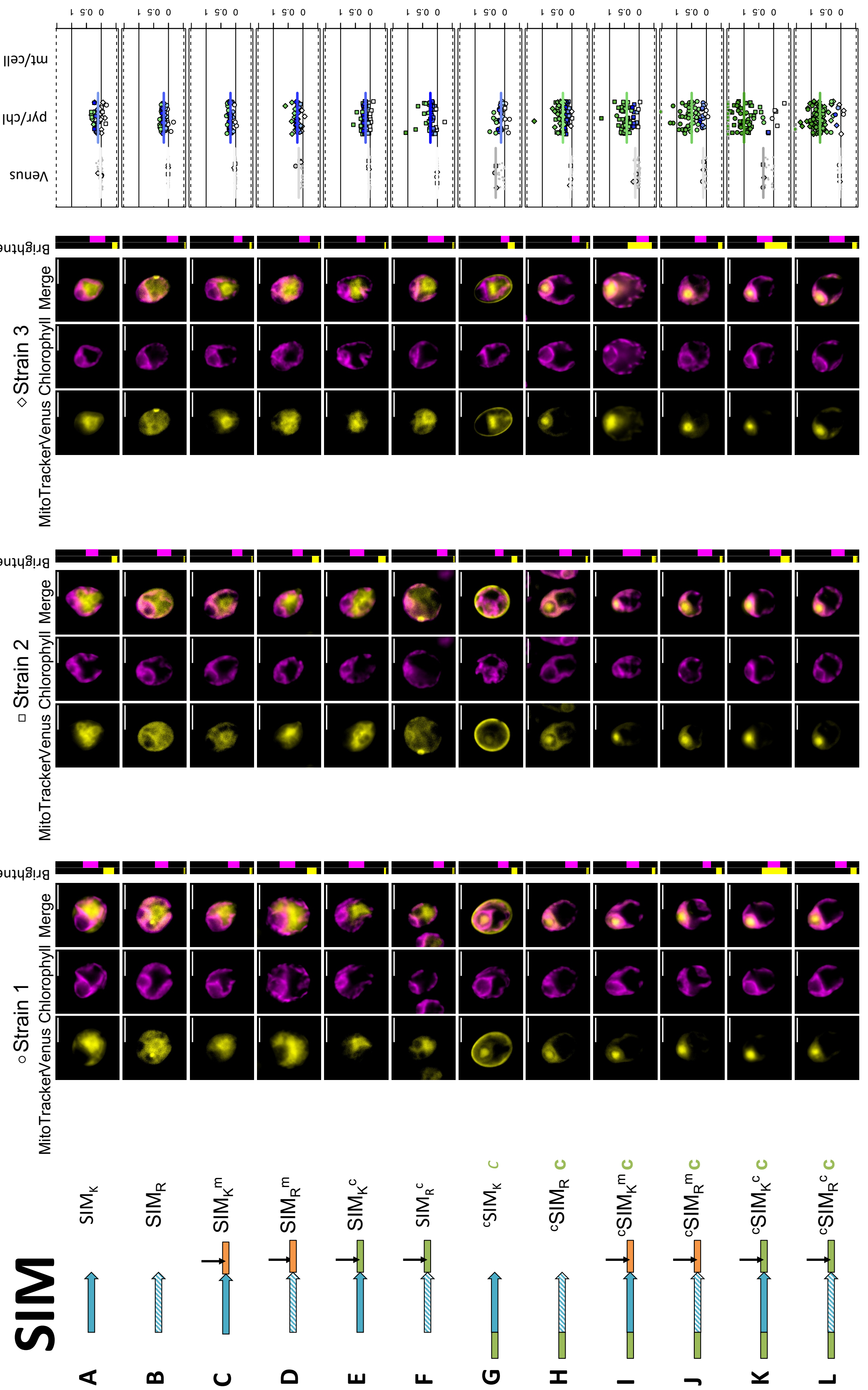


**Fig. S13. Biological replicates of SI Moricin.**

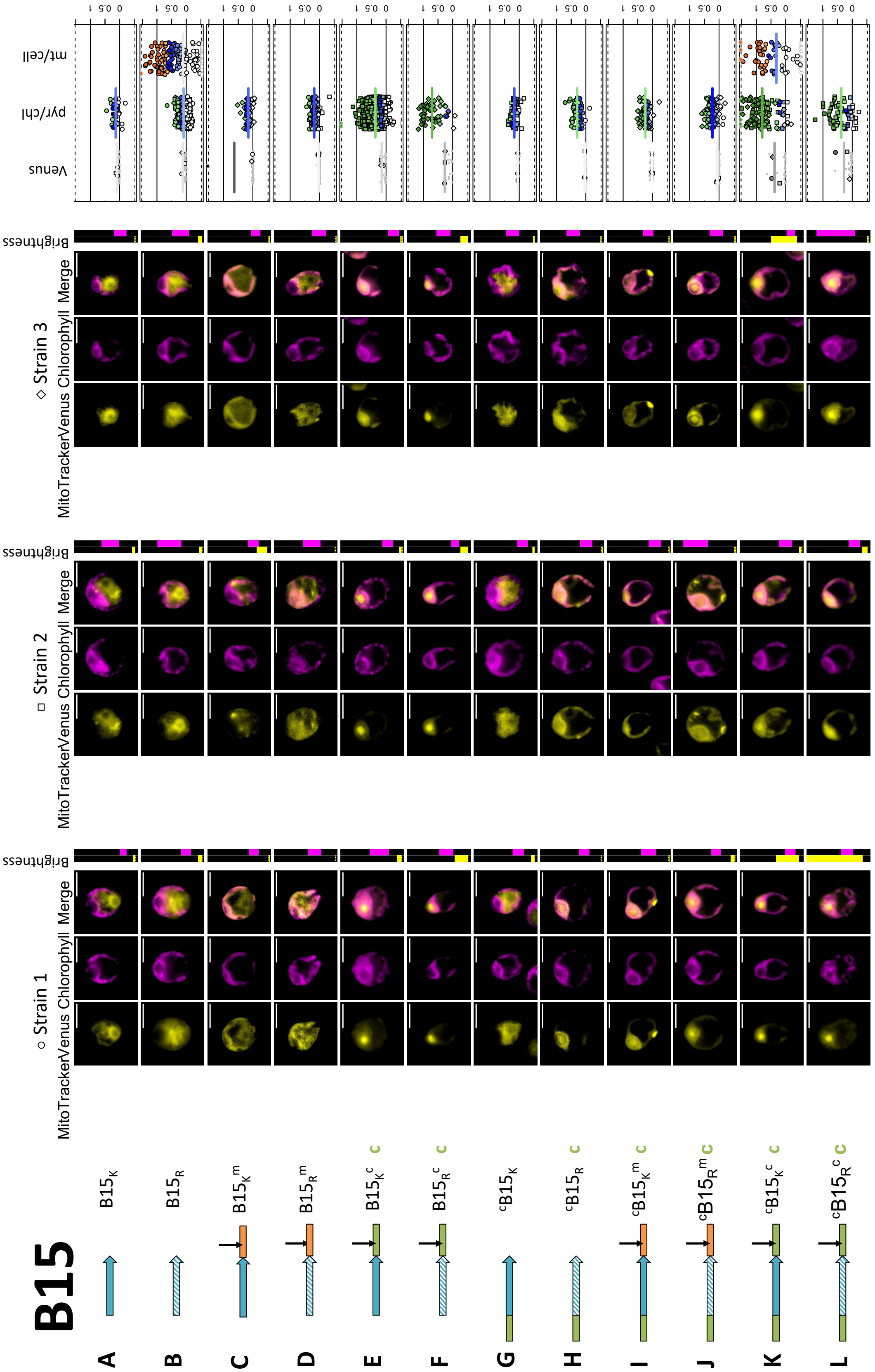


**Fig. S14. Biological replicates of Bacillocin 1580.**

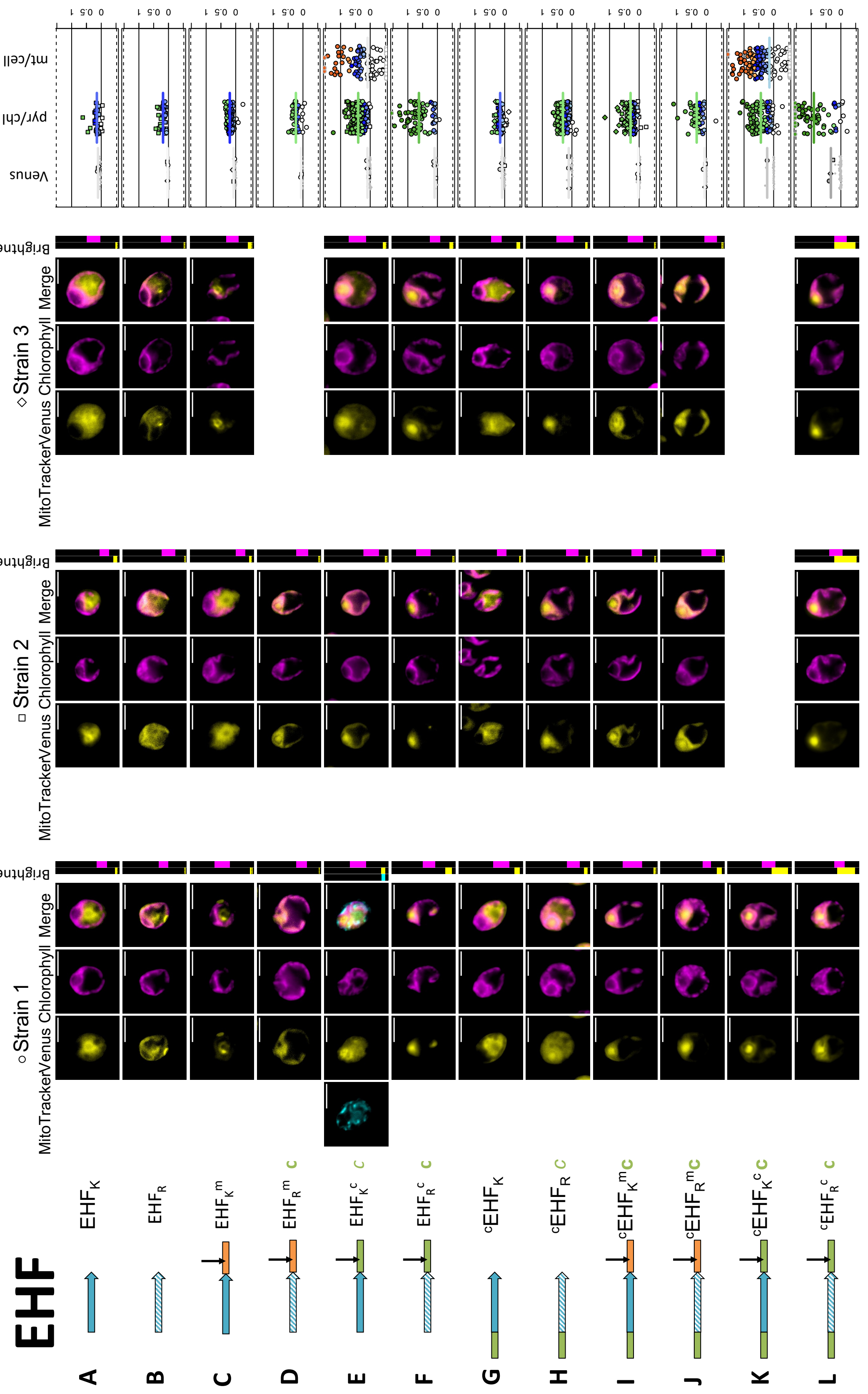


**Fig. S15. Biological replicates of Enterocin HF.**

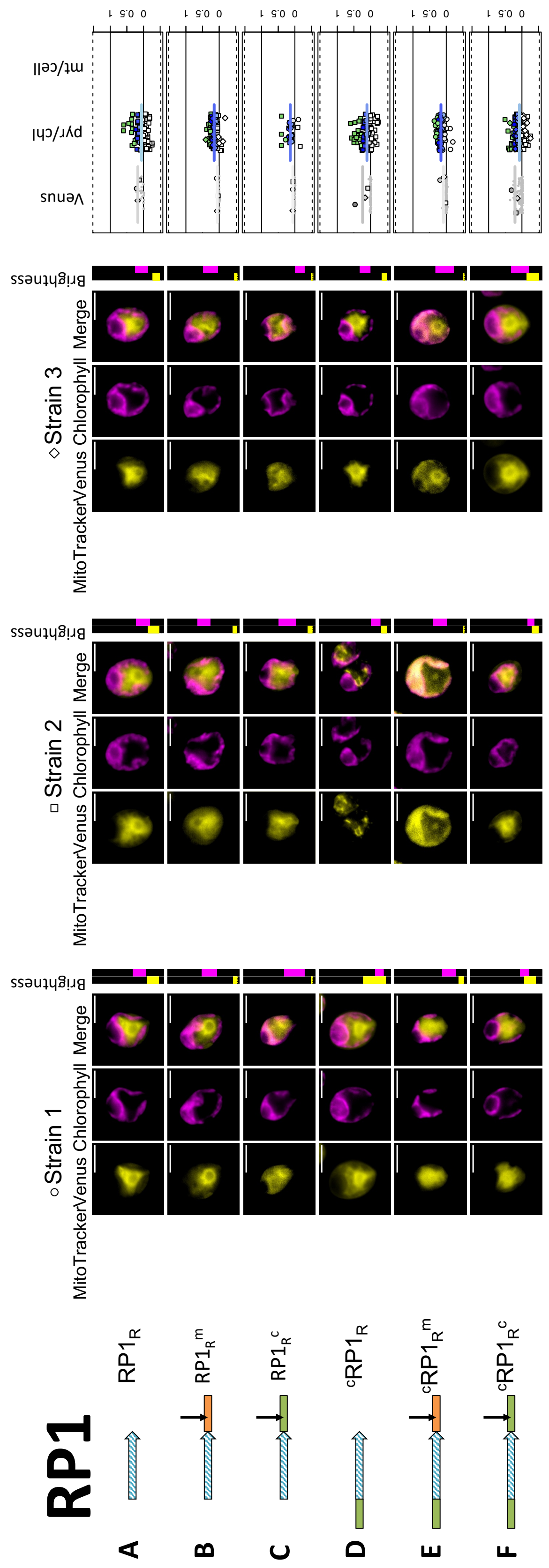


**Fig. S16. Biological replicates of negative control Random Peptide 1.**

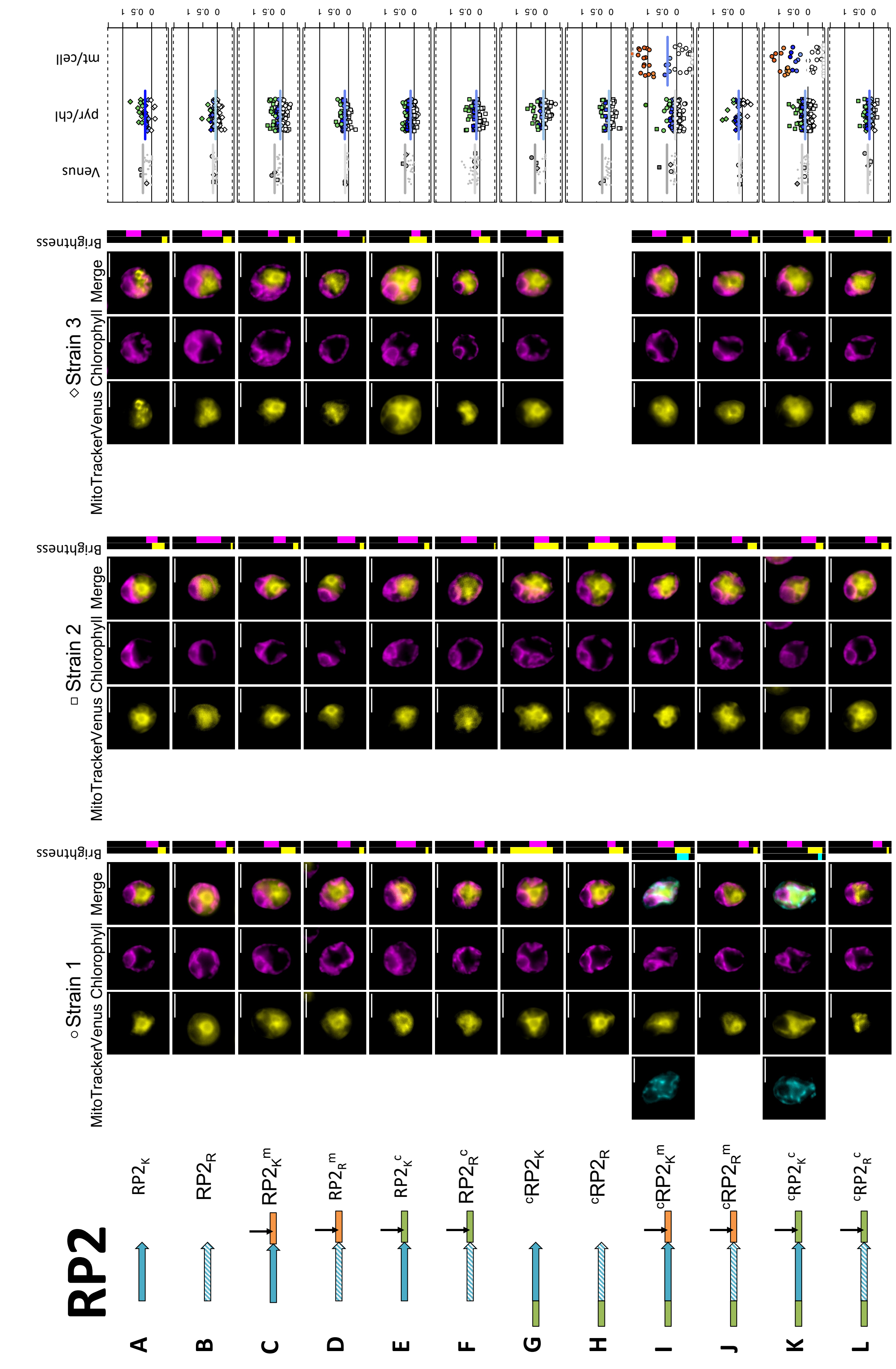


**Fig. S17. Biological replicates of negative control Random Peptide 2.**

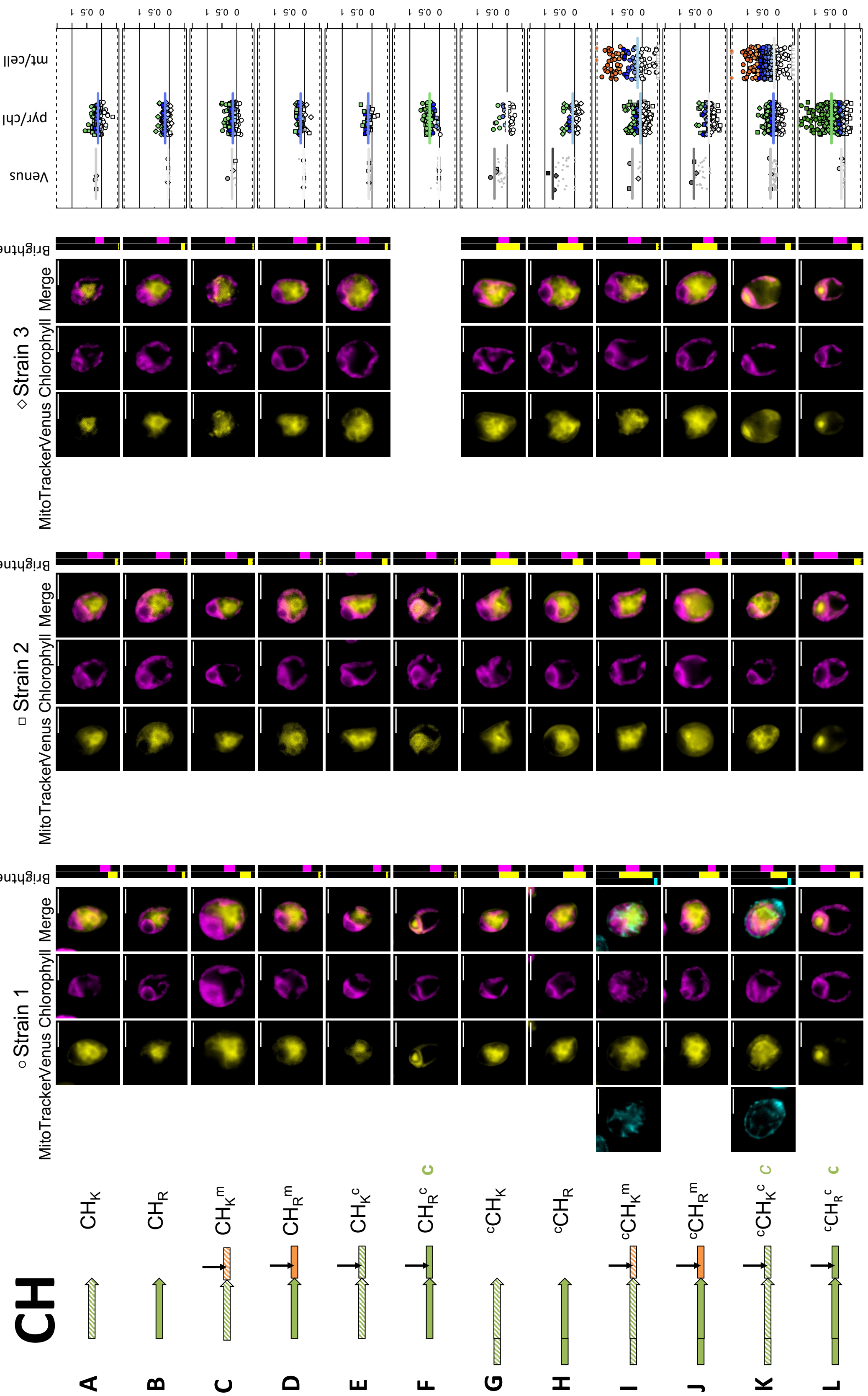


**Fig. S18. Biological replicates of Rubisco activase cTP helical element (CH) control**

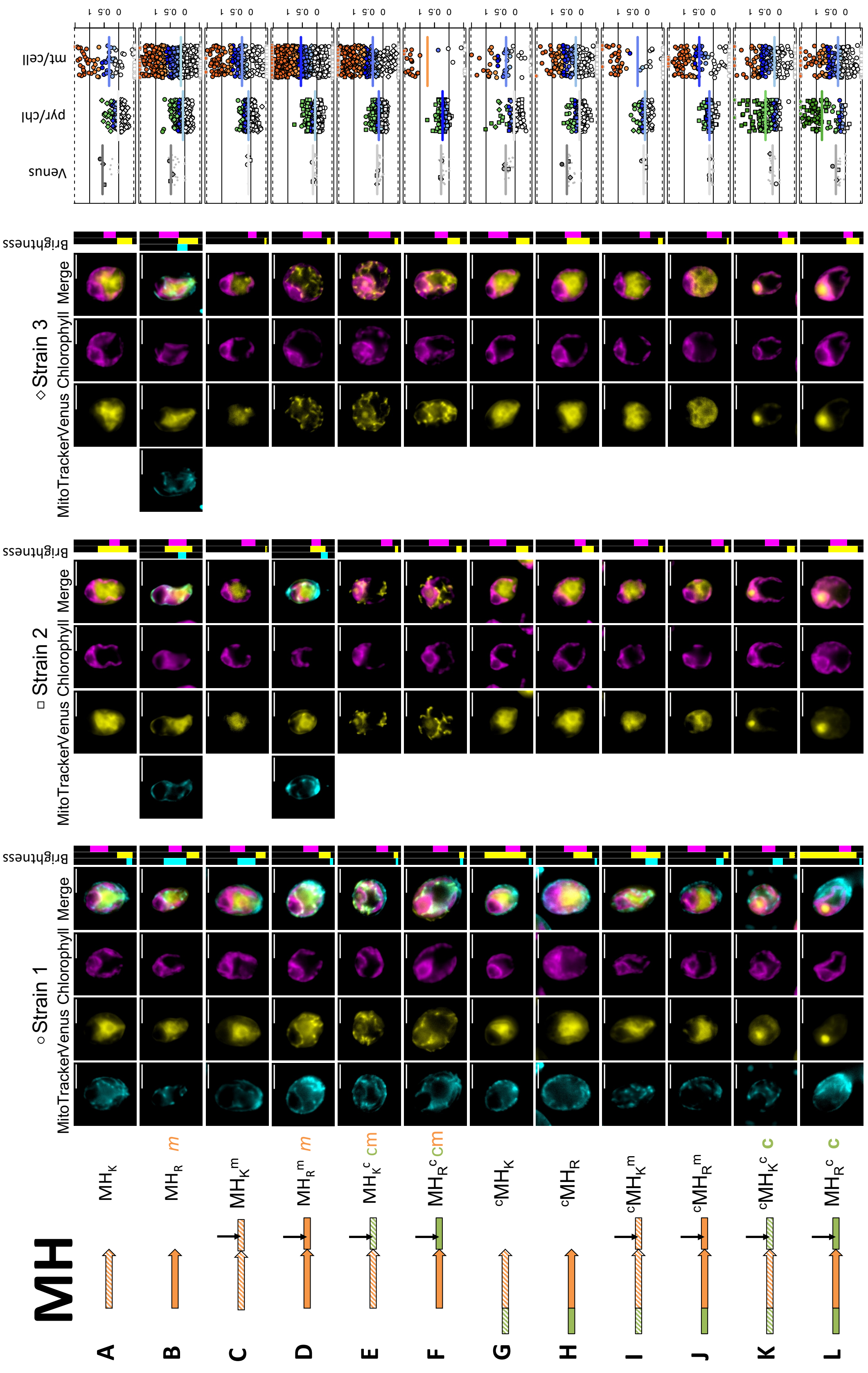


**Fig. S19. Biological replicates of γ-carbonic anhydrase 2 mTP helical element (MH) control**

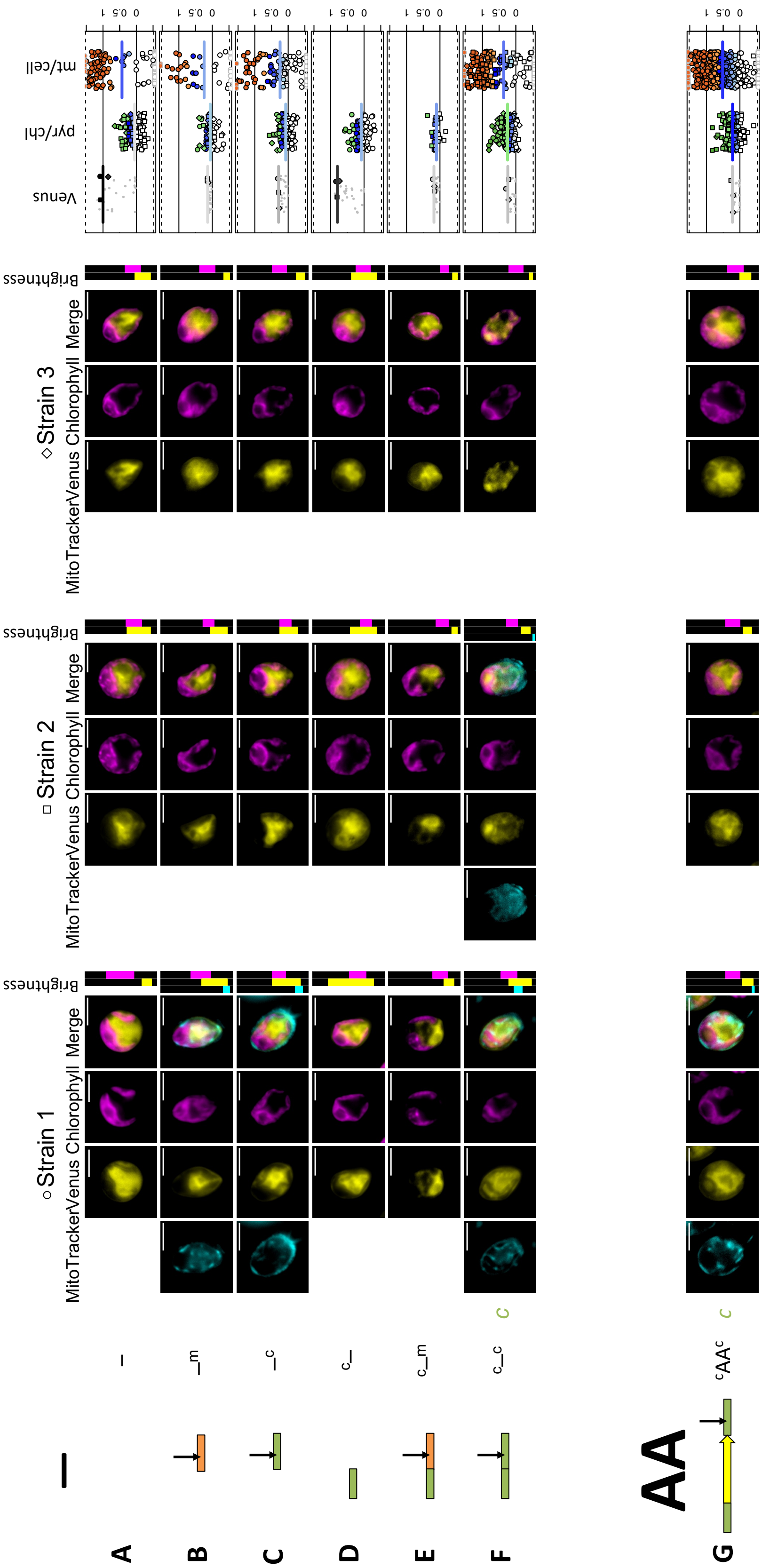


**Fig. S20. Biological replicates of no-peptide and Alanine-screen controls.**

**Fig. S21. Comparison of amino acid frequencies reveals K/R shift.** (**a**) Amino acid frequencies are shown as boxplots (center line: median; box limits: upper and lower quartiles; whiskers: min/max values within 1.5x interquartile range) for Chlamydomonas cTPs in green, Chlamydomonas mTPs in orange and HA-RAMPs in blue. To give a baseline for comparison, the average across UNIPROT is given for each amino acid as red horizontal line. For each residue, different letters underneath distributions indicate a significant difference: groups that share the same letter are significantly different at $p<0.05$ (Multiple Kruskal Wallis tests followed by Dunn post-hoc tests) from groups attributed a different letter. (**b**) Amino acid frequencies are shown as heatmap for human, plant, algal and yeast TPs.

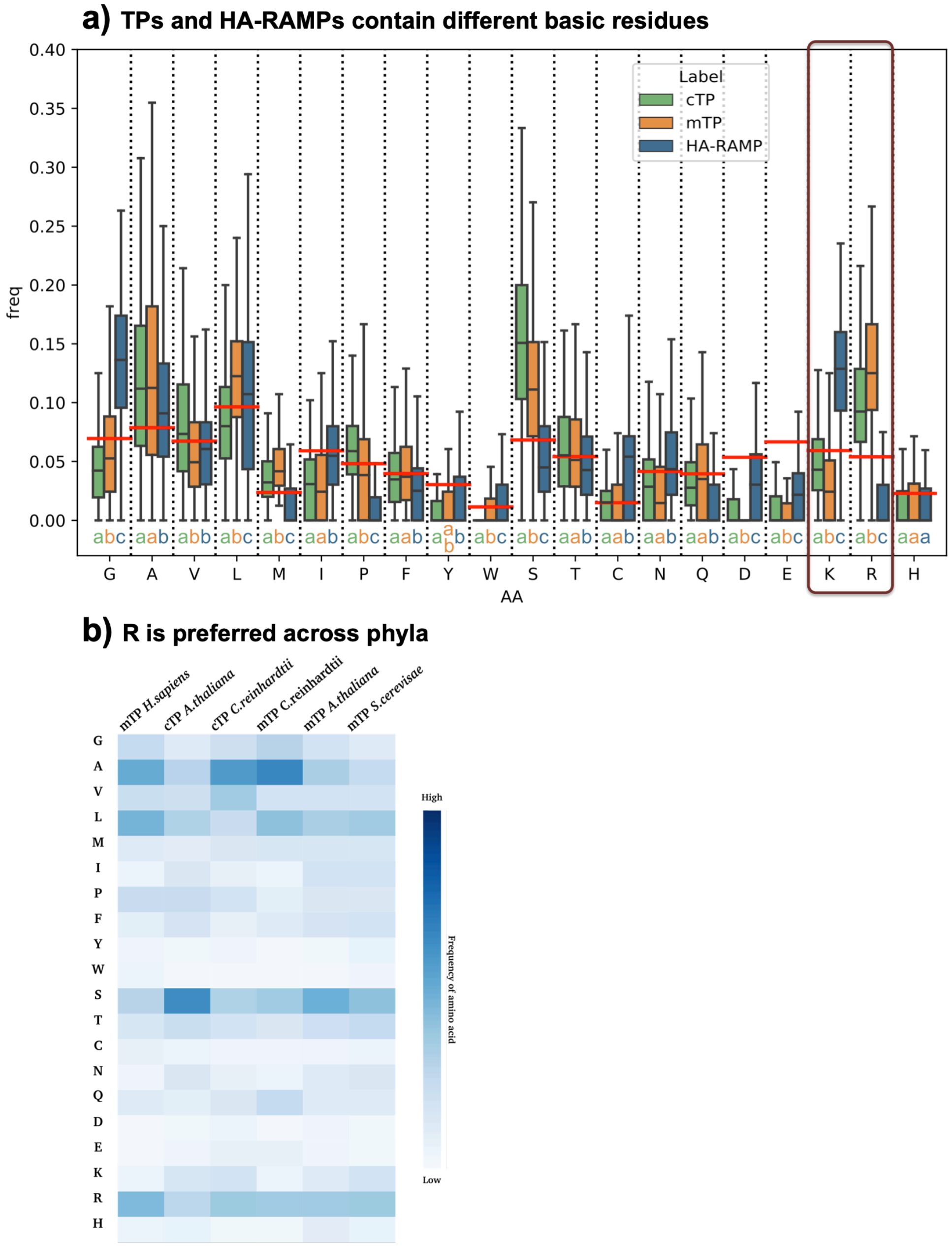

**Fig. S22. Principal component analyses reveal that N- but not C-termini differ between cTPs and mTPs.** Principal component (PC) analyses of auto-cross-correlated Z-scale values (Garrido et al., 2020) for (**a**) N-termini (15 residues) and (**b**) C-termini (33 residues, encompassing -10 to +23 relative to the cleavage site for TPs) respectively of Chlamydomonas cTPs (in green) and mTPs (in orange) as well as the 13 HA-RAMPs under study (in blue; crosses denote HA-RAMPs after K→R).

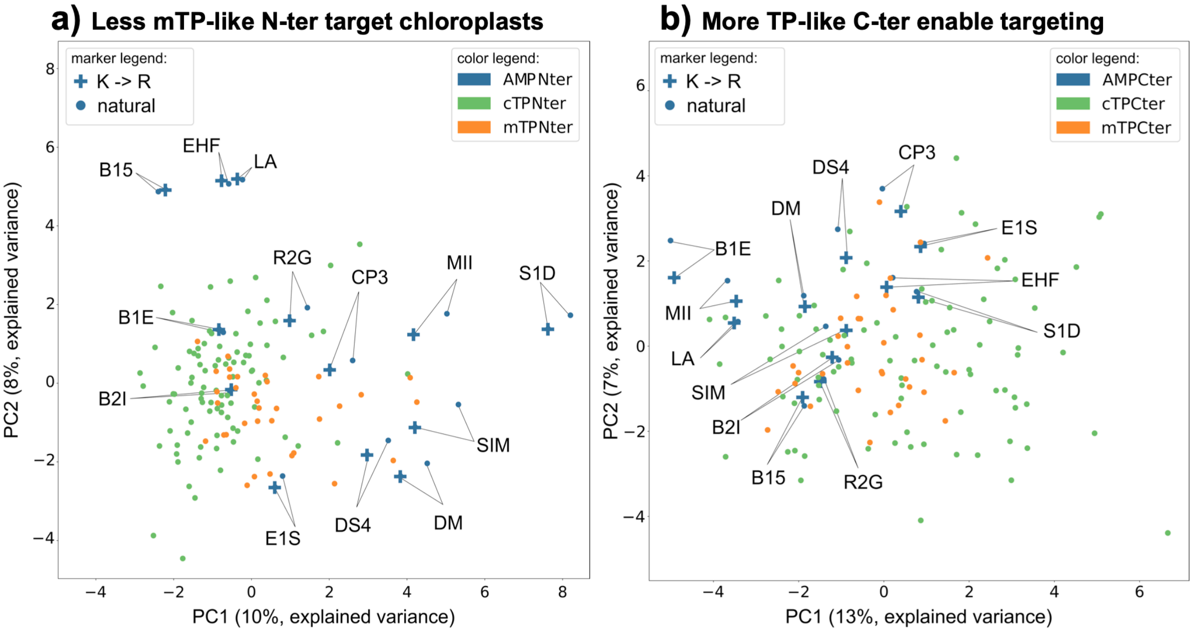

**Fig. S23. Unusual algal cTP N-termini share physicochemical differences against mTP N-termini with plant counterparts.** (**a**) Plots detailing the percentage of uncharged residues (i.e. excluding K/R and D/E, as in (Chotewutmontri et al., 2012) for window sizes ranging from 5 to 17 residues (refer to legend for colour code) show that Chlamydomonas cTPs (on the left) are less uncharged at the N-terminus than Arabidopsis cTPs (on the right)**.** Dashed grey lines indicate the average value for randomized sequences (including mature proteins), to provide an estimate of what would be expected by chance. (**b**) Equivalent graphs (as in A) for mTPs show a similar charge profile across Chlamydomonas and Arabidopsis. (**c-e**) Distributions of salient values as boxplots (center line: median; box limits: upper and lower quartiles; whiskers: min/max values within 1.5x interquartile range) for mTPs in orange and cTPs in green with *Chlamydomonas reinhardtii* (Cr) on the left and *Arabidopsis thaliana* (At) on the right show that: (**c**) Chlamydomonas cTP N-termini contain R at almost the same frequency as mTP N-termini, whereas Arabidopsis cTP N-termini contain fewer R; (**d**) Chlamydomonas TP N-termini are less hydrophobic than Arabidopsis TP N-termini, with plant cTP N-termini showing the highest hydrophobicity; (**e**) Chlamydomonas cTP N-termini are more disordered than mTP or Arabidopsis cTP N-termini. Points represent individual peptides (note that for integer values, point positions are randomized within ±0.5 in y as well as in x to increase point separation), population means are shown as black diamonds. (**f**) Binomial logistic regression classifier separating TPs using auto-cross-covariance of Z-scales (Garrido et al., 2020) of the N-terminal 15 residues for *C. reinhardtii*. The left-hand graph shows the distribution of cTPs in green and mTPs in orange over the model output 'mTP score'. Black bars at the bottom of the graph represent scores for cTPs of Arabidopsis, showing that plant cTPs are recognized as cTPs by the model trained on algal cTP N-termini (89% recognized as cTPs, with an mTP score <0.5). Values for our 13 HA-RAMPs are given below the graph. The right-hand graph shows receiver operating characteristic (ROC) curves, plotting the true positive rate (TPR) against the false positive rate (FPR), where the area under the curve (AUC) serves as estimate of the model quality with values above 0.5 indicating that the model is better than random. (**g**) Equivalent model (as in F) trained on *A. thaliana* TPs, with black bars now representing Chlamydomonas cTPs showing that the majority of algal cTPs are recognized as cTPs by the plant model (75% recognized as cTPs, with an mTP score <0.5).

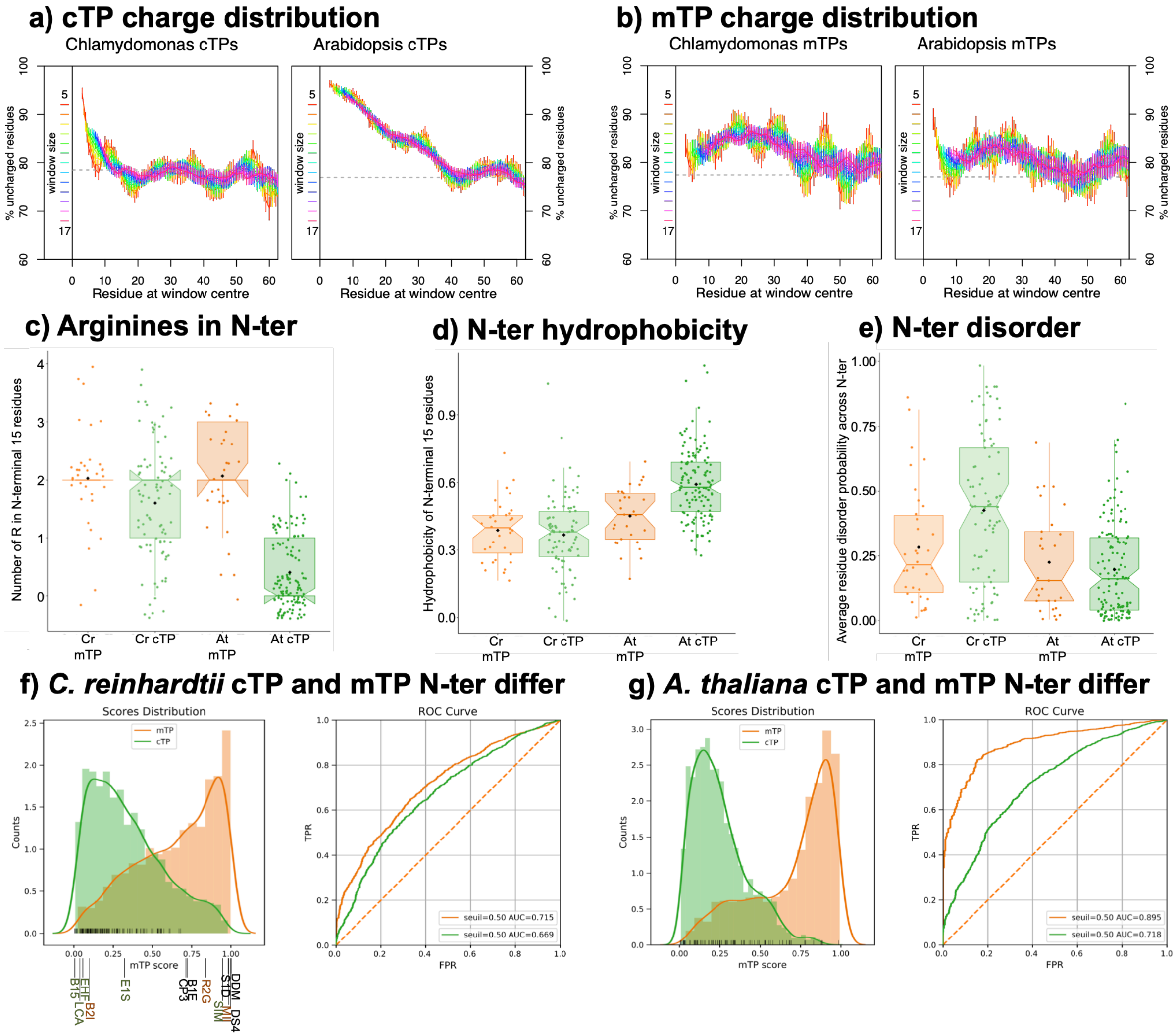
a) cTP charge distribution
Chlamydomonas cTPs
Arabidopsis cTPs
b) mTP charge distribution
Chlamydomonas mTPs
Arabidopsis mTPs
% uncharged residues
window size
Residue at window centre
c) Arginines in N-ter
Number of R in N-terminal 15 residues
Cr mTP
Cr cTP
At mTP
At cTP
d) N-ter hydrophobicity
Hydrophobicity of N-terminal 15 residues
e) N-ter disorder
Average residue disorder probability across N-ter
f) C. reinhardtii cTP and mTP N-ter differ
Scores Distribution
ROC Curve
mTP
cTP
Counts
mTP score
TPR
FPR
seuil=0.50 AUC=0.715
seuil=0.50 AUC=0.669
g) A. thaliana cTP and mTP N-ter differ
seuil=0.50 AUC=0.895
seuil=0.50 AUC=0.718

**Fig. S24. K→R generally improves HA-RAMP targeting.** (**a-e**) Epifluorescence microscopy images of selected examples are shown as in Fig. 3.

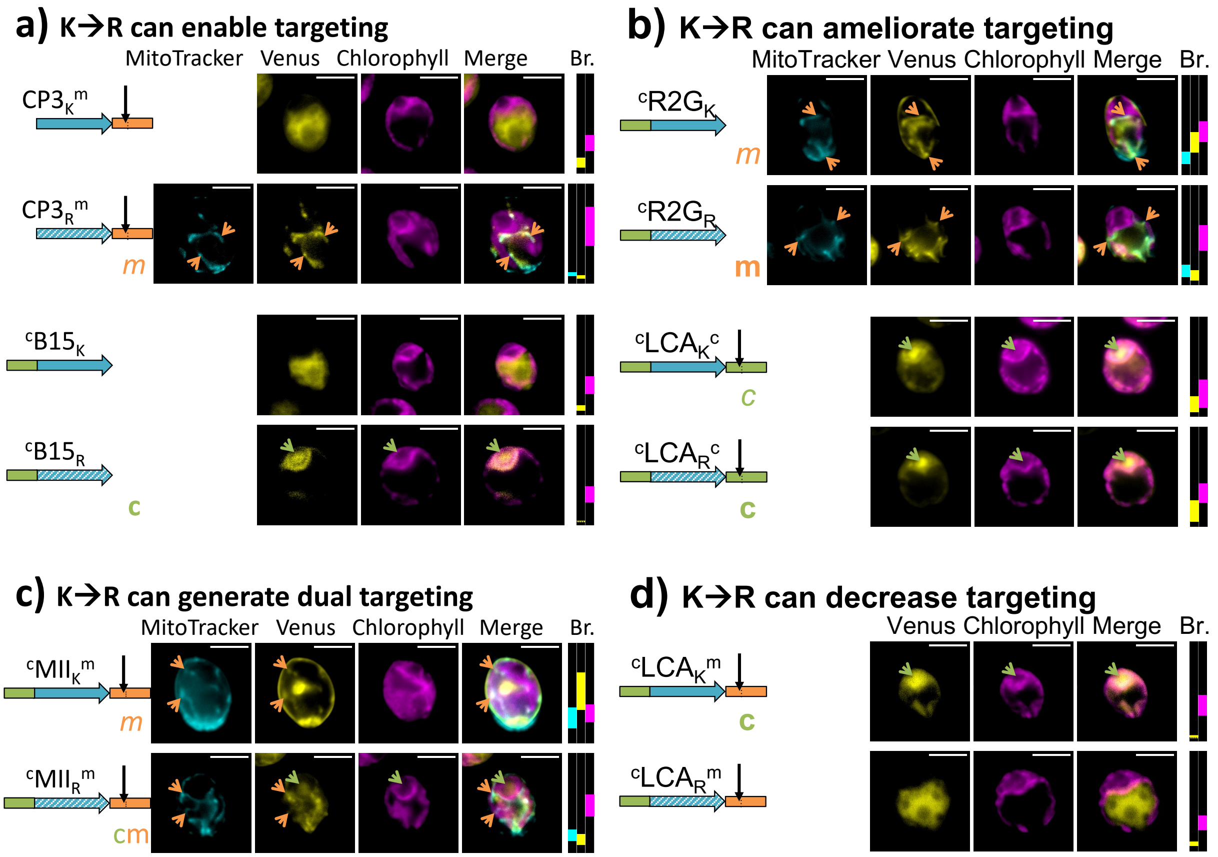

**Fig. S25. HA-RAMP properties determine their targeting propensities.** PCA analysis of cTPs (green dots), mTPs (orange dots), cp-set HA-RAMPs (green crosses), mt-set HA-RAMPs (orange crosses) and other HA-RAMPs (dark crosses) based on their length (Peptide Length: PL), number of residues before the longest predicted amphipathic helix (Long Helix Start: LHS), fraction of residues forming the predicted amphipathic helix (Helix Fraction: HF) and fraction of K (Fraction of Lysines: FK) and R residues (Fraction of Arginine: FR). Arrows on the graph represent the eigenvalues of the individual variables. The first principal component explains 43% of the variance and distinguishes peptides principally according to FK, PL, LHS (pointing left) and FR residues (pointing right). The second principal component explains 24% of the variance and distinguishes peptides principally according to HF. The cp-set HA-RAMPs group with the most distinguishable cTPs, being longer, with a higher LHS and a lower HF compared to mt-set HA-RAMPs and mTPs.

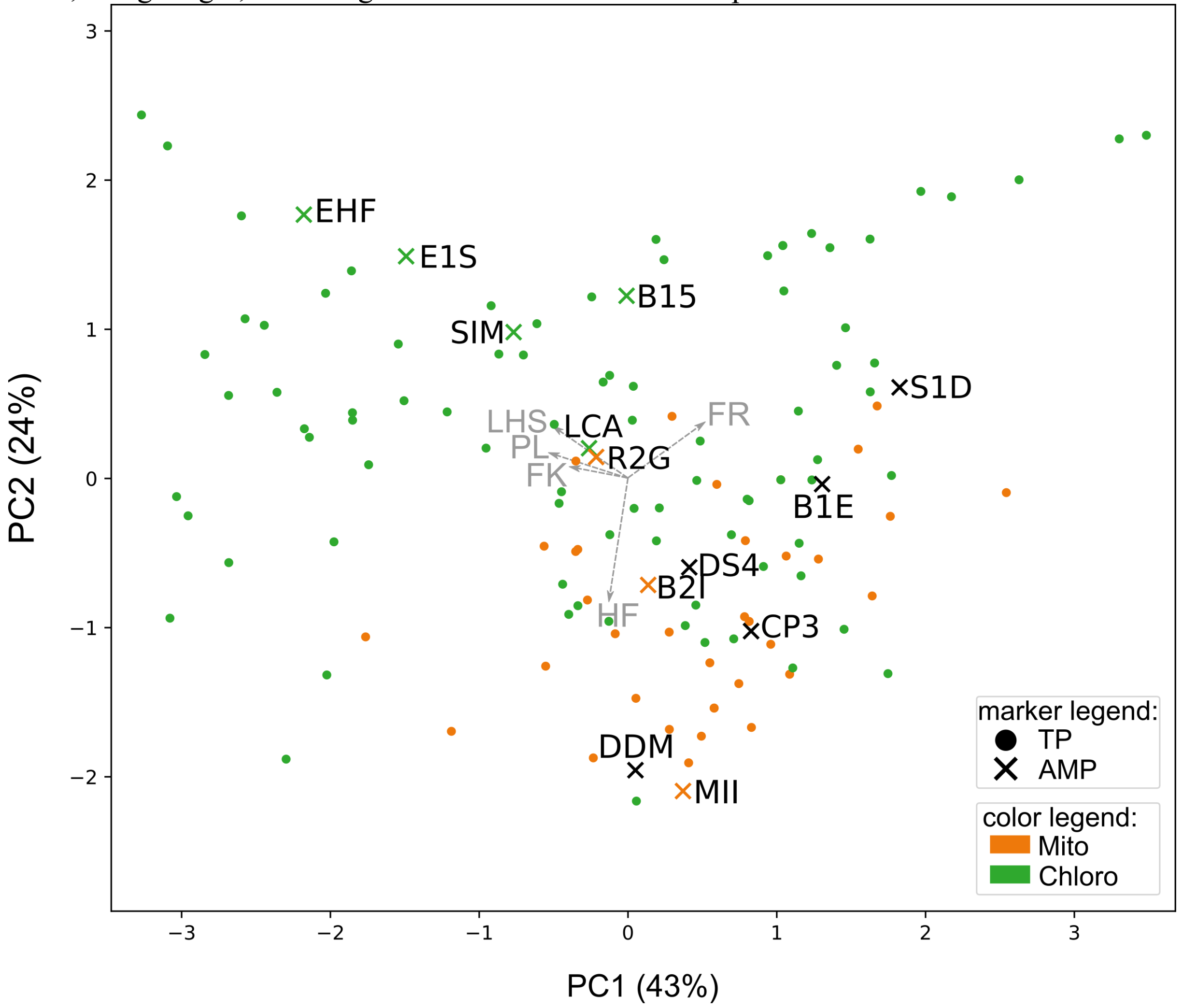

**Fig. S26. Higher protein interactivity predicted for cTPs than mTPs.** (**a,b,f**) For salient properties, Chlamydomonas mTPs and cTPs are compared to HA-RAMPs (cp-set in green, mt-set in orange). Distributions are shown as boxplots (center line: median; box limits: upper and lower quartiles; whiskers: min/max values within 1.5x interquartile range), coloured points represent individual peptides (**a**) cTPs show increased protein interactivity (p=0.0003), as do cp-set HA-RAMPs (p= 0.0355), as estimated through Boman values (Boman, 2003), a proxy developed for AMPs where a value of ca. >2 indicates increased protein interaction potential, and <2 points to membrane interaction. (**b**) cTPs show increased protein interactivity (p<0.0001), and cp-set HA-RAMPs show a similar tendency (p=0.0638), as estimated with Anchor2 (Mészáros et al., 2009), developed to predict the protein interaction potential of disordered sequences. (**c**) Both cTPs and mTPs show a high fraction of peptides (~50%) that have an Hsp70 interaction site over a 6-residue window at any given position along the sequence in the top graph, with a peak towards the N-terminus. HA-RAMP constructs shown in the bottom graph also display a peak in predicted Hsp70-binding sites at the N-terminus, independently of construct localisation. The value obtained after randomizing the position of residues is given as dashed grey line to provide an estimate of how often sites would be expected to occur given amino acid frequencies (~50%). Randomization was done over the entire sequence including the cargo protein. (**d**) For Arabidopsis cTPs in the top graph, the percentage of peptides that have a full 'FGLK' site (Chotewutmontri et al., 2012) (black curve; presence of F and G/P and L/V/A and K/R and absence of D/E within an 8-residue window) far exceeds the value obtained after randomizing the position of residues (dashed black line for full 'FGLK' sites), a proxy for the frequency of the motif expected at random, for most positions along the sequence up to ~60 residues. Randomization was done over the entire sequence including the cargo protein. Reduced 'FGLK'-1 sites containing three out of four elements appear to occur mostly in the context of full FGLK sites, except for 'GLK' sites for which randomization is also shown (dashed red line). Arabidopsis mTPs (bottom graph) also contain 'FLGK' sites at a frequency higher than expected at random between residues 20-40, and 'GLK' sites upstream of residue 40, but motifs are less prevalent than in cTPs. (**e**) Equivalent graphs (as D) for Chlamydomonas show that full 'FGLK' sites (black lines) are rare in cTPs (top graph) and actually more common in mTPs (bottom graph) upstream of residue 40. Reduced 'GLK' sites (red lines) by contrast far exceed the prevalence expected at random (dashed red line) ca. up to residue 40. (**f**) Protein interactivity, estimated through Boman values, for only those parts of peptides that correspond to FGLK-1 sites is increased for cTPs (p<0.0001), and cp-set HA-RAMPs show a similar tendency (p= 0.0627). Note that cTP-C and cTP-N, but not mTP-C, contain 'FGLK-1' sites with predicted protein interactivity. *DS4 contains no 'FGLK-1' sites. Reported p-values were obtained through two-way t-tests for TPs and one-way t-tests for HA-RAMPs.

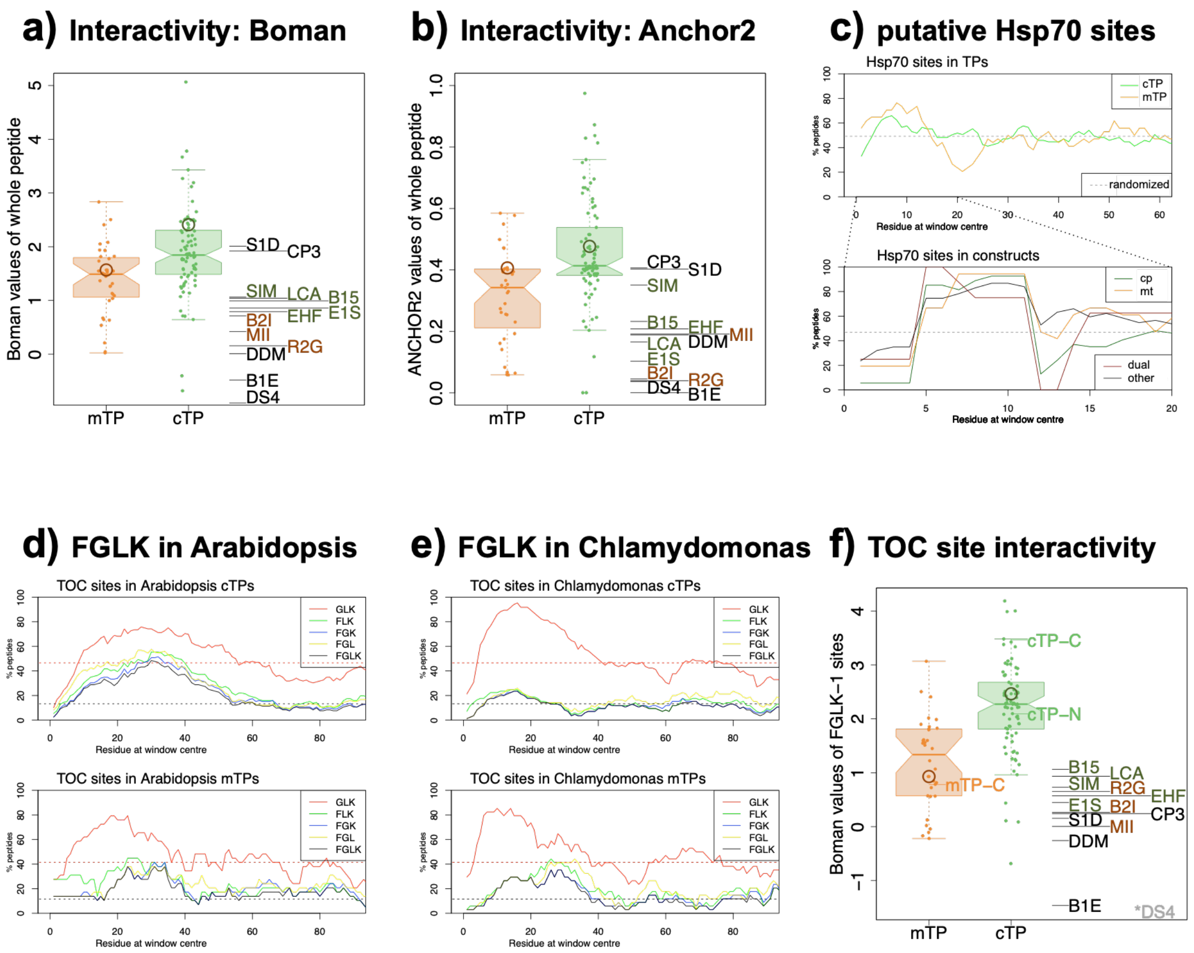

a) Interactivity: Boman
Boman values of whole peptide
mTP
cTP
S1D
CP3
SIM
LCA
B15
EHF
E1S
B2I
MII
R2G
DDM
B1E
DS4
b) Interactivity: Anchor2
ANCHOR2 values of whole peptide
c) putative Hsp70 sites
Hsp70 sites in TPs
randomized
Residue at window centre
Hsp70 sites in constructs
cp
mt
dual
other
% peptides
d) FGLK in Arabidopsis
TOC sites in Arabidopsis cTPs
TOC sites in Arabidopsis mTPs
GLK
FLK
FGK
FGL
FGLK
e) FGLK in Chlamydomonas
TOC sites in Chlamydomonas cTPs
TOC sites in Chlamydomonas mTPs
f) TOC site interactivity
Boman values of FGLK−1 sites
cTP−C
cTP−N
mTP−C
*DS4

**Fig. S27. Hsp70 and (F)GLK sites are present in HA-RAMPs.** Predicted Hsp70 binding sites are underlined in blue. Putative TOC interaction sites are overlined in green: 'FGLK' sites in dark green and 'GLK' sites in light green. See Fig. 1 for detailed annotations. Notably, both TOC and Hsp70-interaction sites are contained within native HA-RAMPs (**a**) and within our TP controls (**b**).

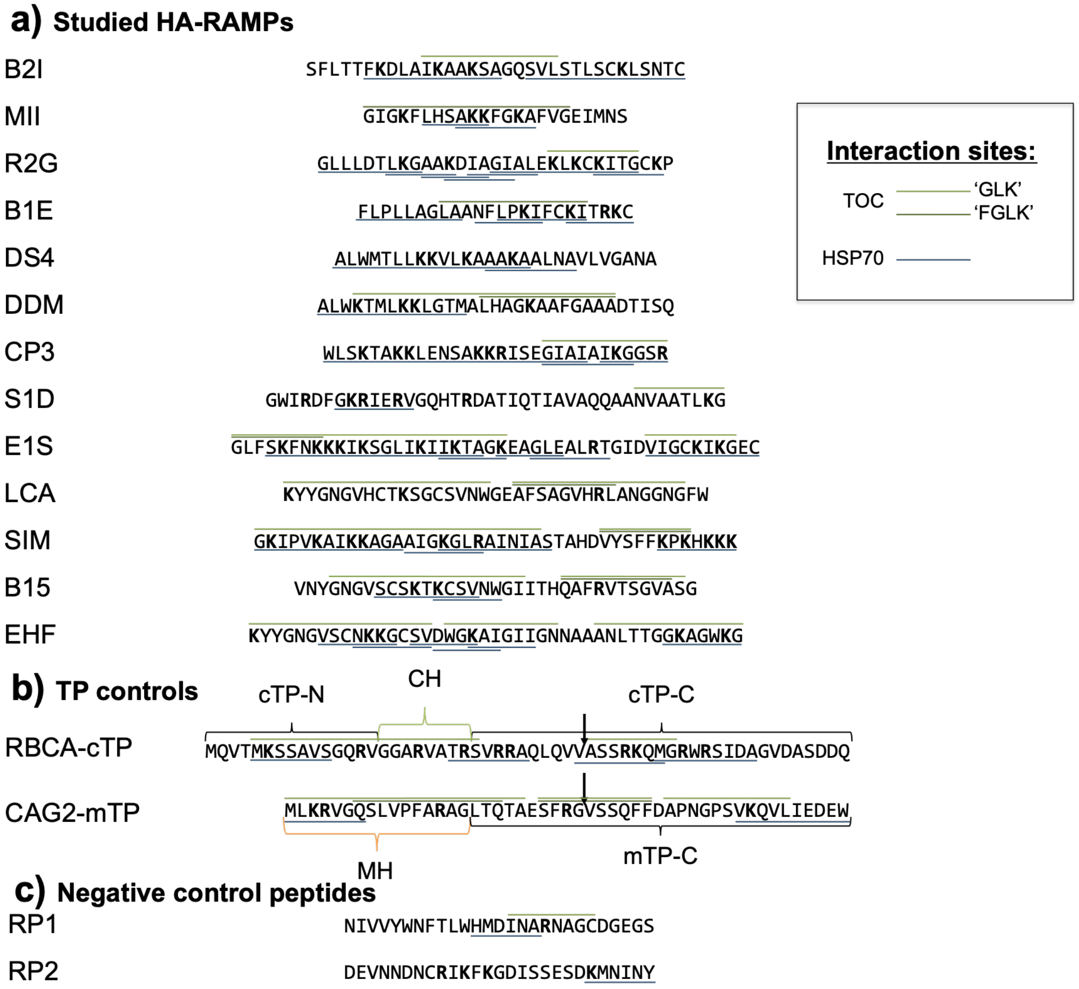

**Fig. S28. Western Blots for K-bearing constructs.** Samples were immunolabelled using an α-FLAG primary antibody. Constructs representing different combinations of TP-element additions are shown in rows A-F, as indicated by a shorthand description and a cartoon (cf. Fig. 2). HA-RAMPs are consistently arranged in lanes a-m. Note that the order of HA-RAMPs differs from the one in Fig. 2. Control constructs in lanes o-r carry the R→K modification. Additional technical controls are present in lanes labeled with greek letters. Sections containing HA-RAMPs, control constructs and technical controls are separated by dotted black lines. Technical controls (cf. Fig. 1): – is no peptide, ${}^{c}CH_{R}^{c}$ is Rubisco activase cTP, $MH_{R}^{m}$ is mitochondrial γ-carbonic anhydrase 2 mTP, ${}^{c}$– is Rubisco activase cTP N-terminal element (15 residues), –${}^{c}$ is Rubisco activase cTP C-terminal element (33 residues, -10 to +23 relative to cleavage site), e.v. is empty vector (no Venus expression). Constructs that generated chloroplast and/or mitochondrial localization are marked with a green 'c' and/or an orange 'm' respectively, in bold for full or in italics for partial targeting. Note that row E contains two separate blots for the left-hand section up to lane i and the right-hand section following lane j.

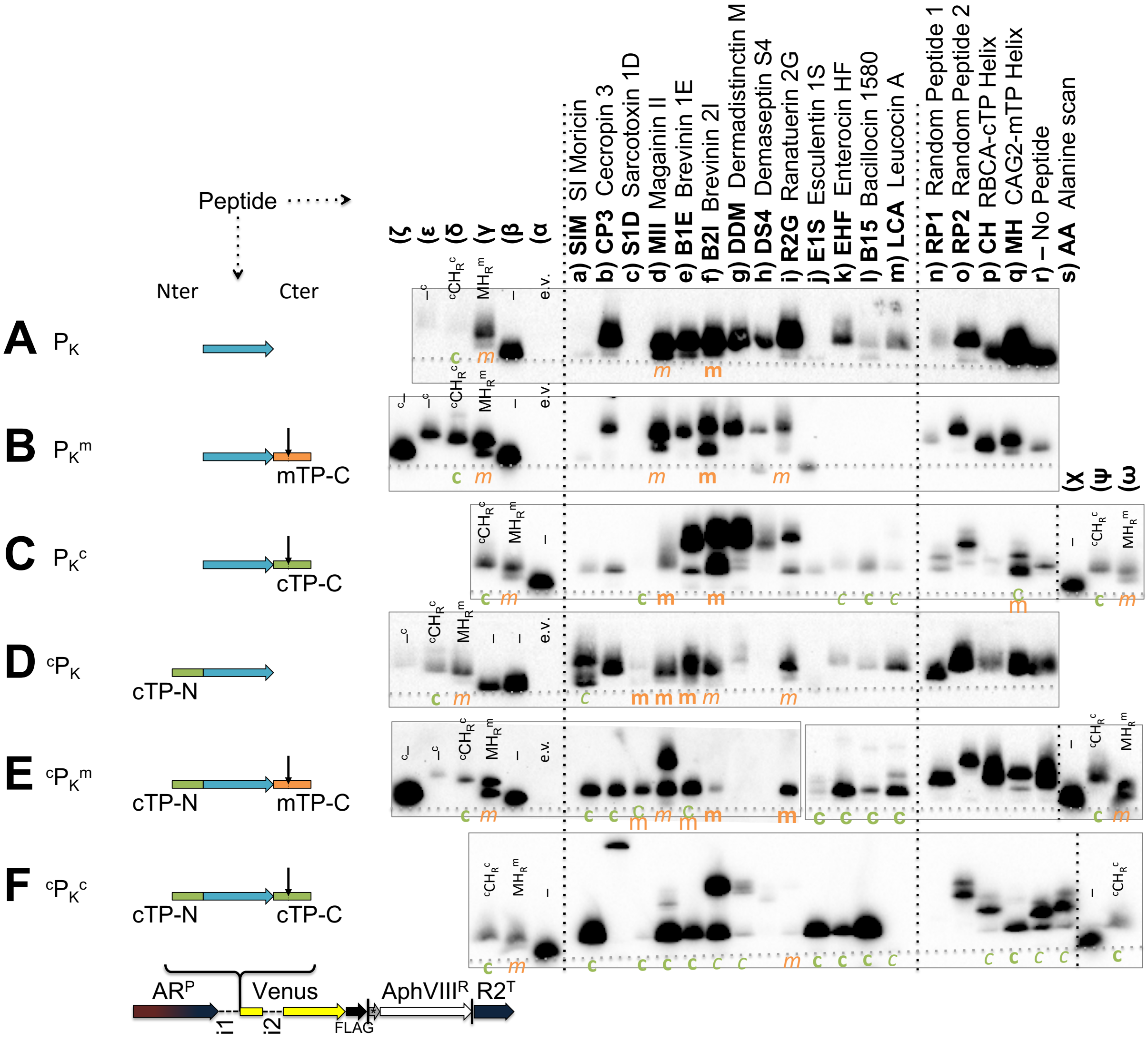
